# From genomic decay to functional advantage: Traits of a persistent, thermally beneficial coral probiotic

**DOI:** 10.64898/2025.12.10.693533

**Authors:** Mei Xie, Congjuan Xu, Nan Xiang, Tianhua Liao, Xing Liu, Zhiqi Liu, Xiaoyuan Feng, Qian He, Zizhen Liang, Weiqi Wang, Yifan Dai, Lili Yan, Claudia Pogoreutz, Lena Barra, Shannon Wing Ngor Au, Liwen Jiang, Christian R. Voolstra, Haiwei Luo

**Author notes:** Corresponding author: Haiwei Luo, School of Life Sciences, The Chinese University of Hong Kong Shatin, Hong Kong SAR.

## Abstract

A key bottleneck in microbiome engineering is ensuring the long-term host association of introduced microbes. Selecting probiotic candidates based on evolutionary genomic decay signatures of emerging host dependency offers a potential solution. The *Ruegeria* strain B4 of the population MC10, identified by such signatures, has shown persistent colonization in corals in a companion study. Whether this persistence translates into a measurable host benefit compared to other coral-associated *Ruegeria* strains, and which mechanisms underlie such benefit, remained unknown. Here we directly compare the probiotic efficacy of MC10-B4 against two sympatric *Ruegeria* strains isolated from the same coral colony and mucus compartment, controlling for host genotype and microenvironment. MC10-B4 inoculation significantly increased heat stress tolerance in the coral model cnidarian Aiptasia (*Exaiptasia diaphana* strain H2), outperforming both sympatric control strains. To understand the mechanistic basis of this probiotic efficacy, we characterized the functional profile of MC10 using an integrated multi-omics approach. The genome of MC10 is enriched in genes associated with host-interaction, including siderophore-mediated iron acquisition and exopolysaccharide biosynthesis, which were phenotypically confirmed by detectable iron scavenging and enhanced biofilm formation. Following exposure to coral tissue extract, MC10-B4 underwent a coordinated motility-to-sessility proteomic reprogramming, downregulating flagellar motor components while upregulating flagellin and biofilm regulators. This response was distinct from its sympatric relatives, which instead mounted broad upregulation of nutrient acquisition systems. MC10-B4’s functional profile, particularly its sensitivity to oxidative stress, contrasts with the traits typically favored in conventional probiotic screens. Our results move beyond the descriptive identification of a coral probiotic by providing initial mechanistic insight into traits associated with long-term host association and improved thermal performance. These findings validate an evolution-guided approach that prioritizes innate colonization potential over pre-defined laboratory functionalities, informing the rational design of durable probiotics.

## Introduction

Probiotic interventions, which administer beneficial microorganisms to improve host health, face a universal bottleneck across medicine, agriculture, aquaculture, and wildlife conservation: the uncertain long-term persistence of introduced microbes within the host [1, 2]. This problem of transient colonization limits the durability of beneficial effects and obscures whether observed improvements stem from transient mechanisms or the establishment of a stable symbiosis. Conventional selection strategies often prioritize predefined functional traits under laboratory conditions, such as antioxidant production or nutrient cycling, without considering colonization success [3]. A paradigm that considers a microbial lineage’s innate potential for stable host association first could address this bottleneck.

An evolution-guided framework for selecting probiotics was proposed to overcome this limitation [4]. This approach inverts the conventional pipeline by first identifying bacterial lineages that exhibit genomic signatures suggestive of an irreversible evolutionary transition toward host dependency, such as insertion sequence expansion and pseudogenization of core metabolic pathways [5]. These signatures are established hallmarks of bacteria adapting to a host-associated lifestyle. The central hypothesis posits that such an evolutionary trajectory intrinsically predisposes bacteria for persistent host colonization. Subsequent screening for beneficial host effects then identifies candidates with combined colonization potential and probiotic function.

Coral reefs represent a critical test case for this framework. These foundational marine ecosystems are experiencing severe global decline due to climate-driven mass bleaching [6–8]. Probiotic therapy is a promising strategy to enhance coral resilience [2], but scalable application is constrained by the transient colonization issue. Application of the evolution-guided framework to a collection of coral-associated bacteria led to the identification of the *Ruegeria* population MC10 as a persistently associated probiotic candidate [4]. Strain MC10-B4, chosen for its pronounced genomic decay, demonstrated sustained retention for the entire 8-month monitoring period in field-outplanted corals that enhanced host thermal resilience during a natural bleaching event. This proof-of-concept validated the framework’s potential to select for persistent probiotics. It also created a knowledge gap because the functional and mechanistic bases underlying MC10’s successful long-term residency and benefit remained uncharacterized.

The companion study established that MC10-B4 persistently associates with its host [4]. Whether this persistence was accompanied by a discernable host benefit, however, remains untested. The present study directly addresses this question through a comparative assessment of probiotic efficacy using standardized thermal tolerance assays in a model cnidarian. Three strains isolated from the same coral colony and mucus compartment were compared: MC10-B4 and two sympatric controls (MC0-A5 and MC15-BG7). This design identifies traits linked specifically to the evolutionary trajectory of MC10 while controlling for host genotype and microenvironment. The experimental design tests two hypotheses. First, MC10’s genomic signatures of host dependency are associated with a distinct suite of realized phenotypic traits, including biofilm formation and iron acquisition, that facilitate stable colonization. Second, MC10 exhibits a host-responsive proteomic reprogramming distinct from non-candidate relatives, reflecting metabolic integration with the host. Genomic predictions were examined through targeted phenotypic assays, and proteomic responses to host metabolites were compared across the three sympatric bacterial strains. By linking genomic signatures to functional profiles and differential probiotic efficacy, this work establishes a mechanistic basis for how evolutionary trajectory can inform the design of stable, effective probiotics.

## Materials and Methods

Coral sampling, bacterial isolation, genome sequencing, and population delineation of 419 *Ruegeria* strains from five coral species across nine Hong Kong reef sites were performed as described in the companion study [4]. In this work, *Ruegeria* populations (termed main clusters, MCs) were delineated using PopCOGenT [9], which identifies genetically cohesive groups based on patterns of homologous recombination. Among the 116 populations identified, MC10 exhibited genomic signatures of emerging host dependency, including insertion sequence expansion and pseudogene proliferation, and was therefore selected as a candidate probiotic. Building on this foundation, we conducted comparative genomic analyses using the 34 closed genomes (15 from MC10, two from its sister MC46, and one each from 17 other MC populations), supplemented with the publicly available genome of the pelagic model *Ruegeria pomeroyi* DSS-3.

To assess probiotic efficacy, we inoculated the cnidarian model Aiptasia H2 (*Exaiptasia diaphana*) [10] with three *Ruegeria* strains: MC10-B4 (a representative strain from the MC10 population), MC0-A5, and MC15-BG7. Host responses were quantified using the Coral Bleaching Automated Stress System (CBASS) [11, 12] (Fig. S1). A 0.22 μm-filtered seawater (FSW) placebo group was included as a baseline. Maximum photosynthetic efficiency (F_v_/F_m_) was measured after 1 h dark acclimation at T7 (7th hour after CBASS assay start, immediately post-stress) and T18 (18th hour after CBASS assay start, after 11 h recovery) using a MINI-PAM-II fluorometer (Walz, Germany) [13]. Thermal tolerance thresholds (ED50) were calculated as the temperature at which F_v_/F_m_ declined by 50% relative to unstressed controls [14].

Orthologous gene families were identified with OrthoFinder v2.5.1 [15]. Prophage regions were predicted using the “What the Phage” workflow [16] and refined with PHASTEST v1.0.0 [17]. Toxin-antitoxin systems were predicted using antiSMASH v7.0 [18]. DMSP degradation-related genes were annotated using the DiTing database v0.95 [19]. KEGG pathway enrichment analysis compared 15 MC10 genomes against 20 non-MC10 *Ruegeria* genomes using phylogenetic generalised least squares (PGLS). Binary Phylogenetic Generalized Linear Mixed Model (BinaryPGLMM) analysis across the 35 closed genomes identified individual genes with differential presence while controlling for shared evolutionary history.

Targeted phenotypic assays compared MC10-B4 against two sympatric control strains (MC0-A5 from population MC0 and MC15-BG7 from population MC15), isolated from the same *O. crispata* colony and mucus compartment in the companion study; the pelagic model strain *R. pomeroyi* DSS-3 was included as a benchmark. These assays examined intracellular inclusions via transmission electron microscopy, motility (swimming, swarming, twitching, and sedimentation), biofilm formation via crystal violet staining, siderophore production via chrome azurol S assay, vitamin prototrophy in defined marine basal medium, oxidative stress tolerance on H_2_O_2_-supplemented agar, and catalase activity by bubble formation.

For proteomic analysis, bacterial cultures at their exponential phase were exposed to macerated coral tissue extract or an autoclaved filtered seawater control for 4 hours. Proteins were separated by SDS-PAGE, digested with trypsin, and analyzed by liquid chromatography-tandem mass spectrometry on an Orbitrap Fusion Lumos Tribrid mass spectrometer. Raw spectra were processed with Proteome Discoverer v3.1.0 [20] and matched against custom strain-specific protein databases generated from genomes sequenced in the companion study. Label-free quantification with retention time alignment and feature mapping determined relative protein abundance. Beta diversity was evaluated via PCoA based on Bray-Curtis dissimilarity. Differential expression analysis employed the limma algorithm with significance thresholds of FDR < 0.05 and |log_2_FC| > 0.5.

Detailed protocols for all methods, including culture conditions, bioinformatic parameters, assay procedures, and statistical analyses are provided in Supplementary Materials.

## Results

### *Ruegeria* MC10 provides superior thermal protection in a model cnidarian

The evolution-guided framework predicted that candidates selected for genomic signatures of early-stage host dependency would exhibit enhanced colonization potential with evidence for an increase in thermal resilience [4]. To confirm this notion, we assessed the presence of a thermal benefit following inoculation of MC10-B4 in comparison to two sympatric control strains MC0-A5 and MC15-BG7. These three strains were isolated from the same *Oulastrea crispata* colony and mucus compartment, thereby controlling for variation attributable to host genotype, micro-environment, and sampling time.

In standardized acute heat-stress assays using the model cnidarian Aiptasia H2, MC10-B4 inoculation significantly increased host thermal tolerance compared to the FSW placebo, with a 1.0°C higher ED50 (thermal tolerance threshold) at T7 (*P* = 0.001) and a 0.6 °C higher ED50 at T18 (*P* < 0.05) (Fig. 1; Dataset S1). MC0-A5 provided an intermediate increase in thermal tolerance, elevating ED50 by 0.6 °C at T7 (*P* < 0.01) but showing no significant increase after recovery (0.3 °C at T18, *P* = 0.13). MC15-BG7 conferred minimal benefit, with a 0.3 °C ED50 increase at T7 (*P* < 0.05) and no detectable effect at T18 (0.02 °C, *P* = 0.97). As such, MC10-B4 was the only strain that significantly improved thermal tolerance after the recovery phase.

**Fig. 1.**
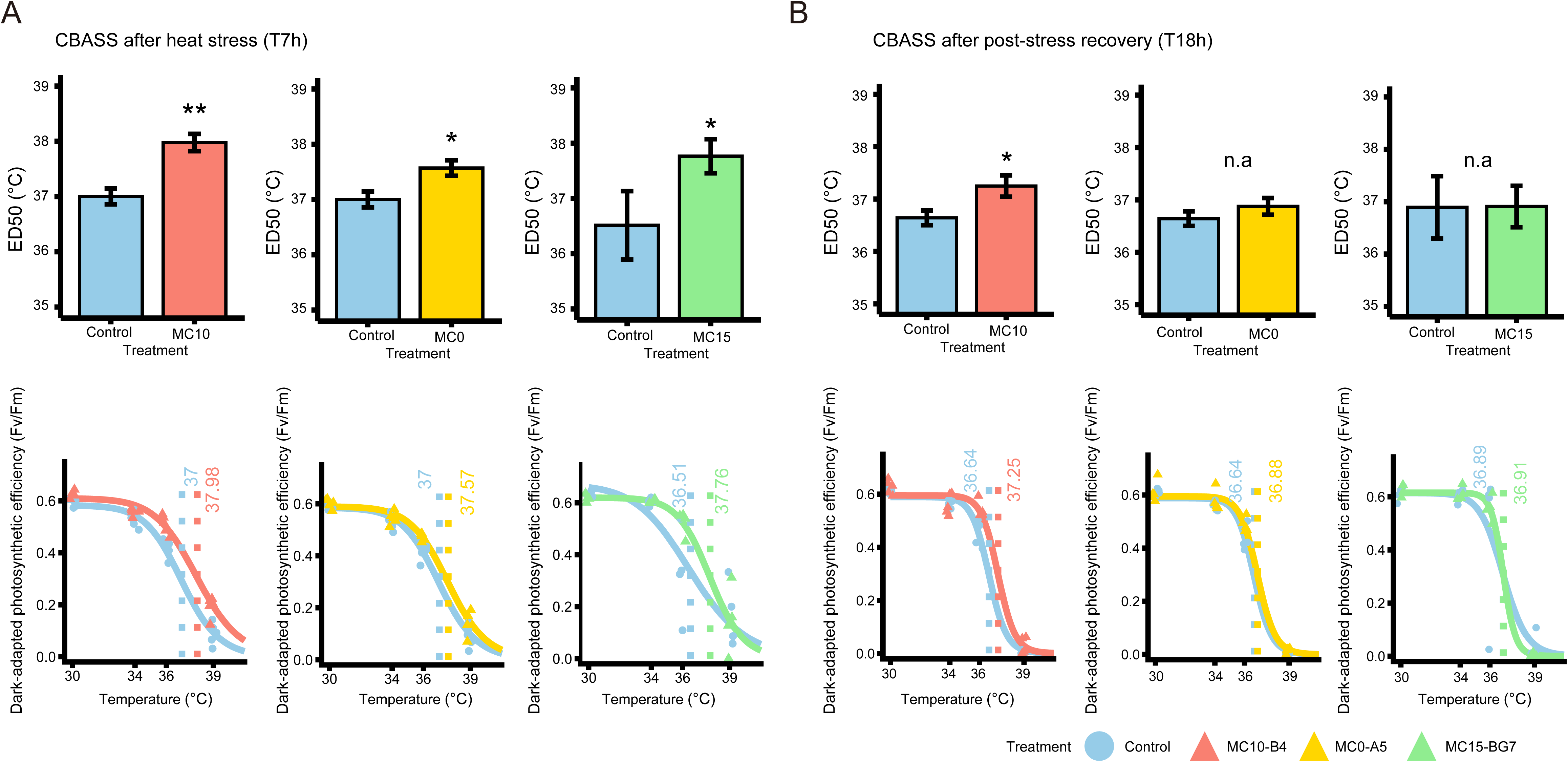
Effect of *Ruegeria* inoculation on *Exaiptasia diaphana* H2 heat tolerance and resilience through CBASS assays. Maximum photosynthetic efficiency (F_v_/F_m_) over temperature curves and determined ED_50_ thermal tolerance thresholds is used as a proxy for bleaching susceptibility of the host inoculated with *Ruegeria* strains MC10-B4, MC0-A5, and MC15-BG7 after (**A**) heat stress (T7) and (**B**) post-stress recovery (T18). In each panel, the upper plots show ED_50_ values indicated as thermal tolerance thresholds, and the lower plots show ED_50_ modeling based on log-logistic regression of F_v_/F_m_ values across temperatures. F_v_/F_m_ was measured for Aiptasia H2 after 1 h of dark acclimation using a MINI-PAM-II fluorometer. Data represented mean ± standard errors (independent Student’s t-tests, *P* < 0.05, *; *P* < 0.01, **; n.a = no significant difference). *Ruegeria* treatment-specific ED_50_ estimates were shown as vertical lines. Blue: FSW control; red: MC10-B4; yellow: MC0-A5; green: MC15-BG7.

### Comparative genomics reveals *Ruegeria* MC10’s unique repertoire of host-adaptation genes

To identify the genetic features underlying MC10’s enhanced colonization and host thermal protection, we characterized its genomic architecture and gene content in comparison to other coral-associated *Ruegeria* lineages. A companion study [4] established the foundation for this work, including isolation of 419 *Ruegeria* strains from five coral species, generation of 35 high-quality closed genomes (15 MC10 plus 20 non-MC10 genomes), and delineation of 116 distinct sub-species populations (Main Clusters, MCs) using PopCOGenT (Fig. S2).

Comparative genomic analysis of the closed genomes revealed that the MC10 population possesses a chromosome averaging 4.52 ± 0.02 Mb (except MC10-AC11 with a less complete assembly at 3.51 Mb), compared to other *Ruegeria* MCs that typically range from 3.22 to 4.04 Mb (Fig. 2B). A defining architectural feature was the integration of plasmid-derived sequences into the chromosome, a phenomenon not observed in other lineages (Fig. 2A). The extrachromosomal plasmids carried by MC10 isolates (ranging from two to three per genome) were exclusively associated with this population and were notably smaller (0.133 to 0.443 Mb) than the large, often >1 Mb plasmids found in other MCs. MC10 genomes also contained more integrated prophage regions, harboring four to five intact prophages (e.g., prophages in MC10-B4 labeled in Dataset S2) spanning 120-160 Kbp per genome, with a cumulative size significantly greater than the 0-3 prophages (totalling <70 Kbp), typical of other lineages (phylANOVA, *P* < 0.05; Fig. 2C, Fig. S3);except for MC45-AM7, which had a comparable profile. These architectural signatures are conserved across all MC10 isolates, indicating they originated in the population’s last common ancestor.

**Fig. 2.**
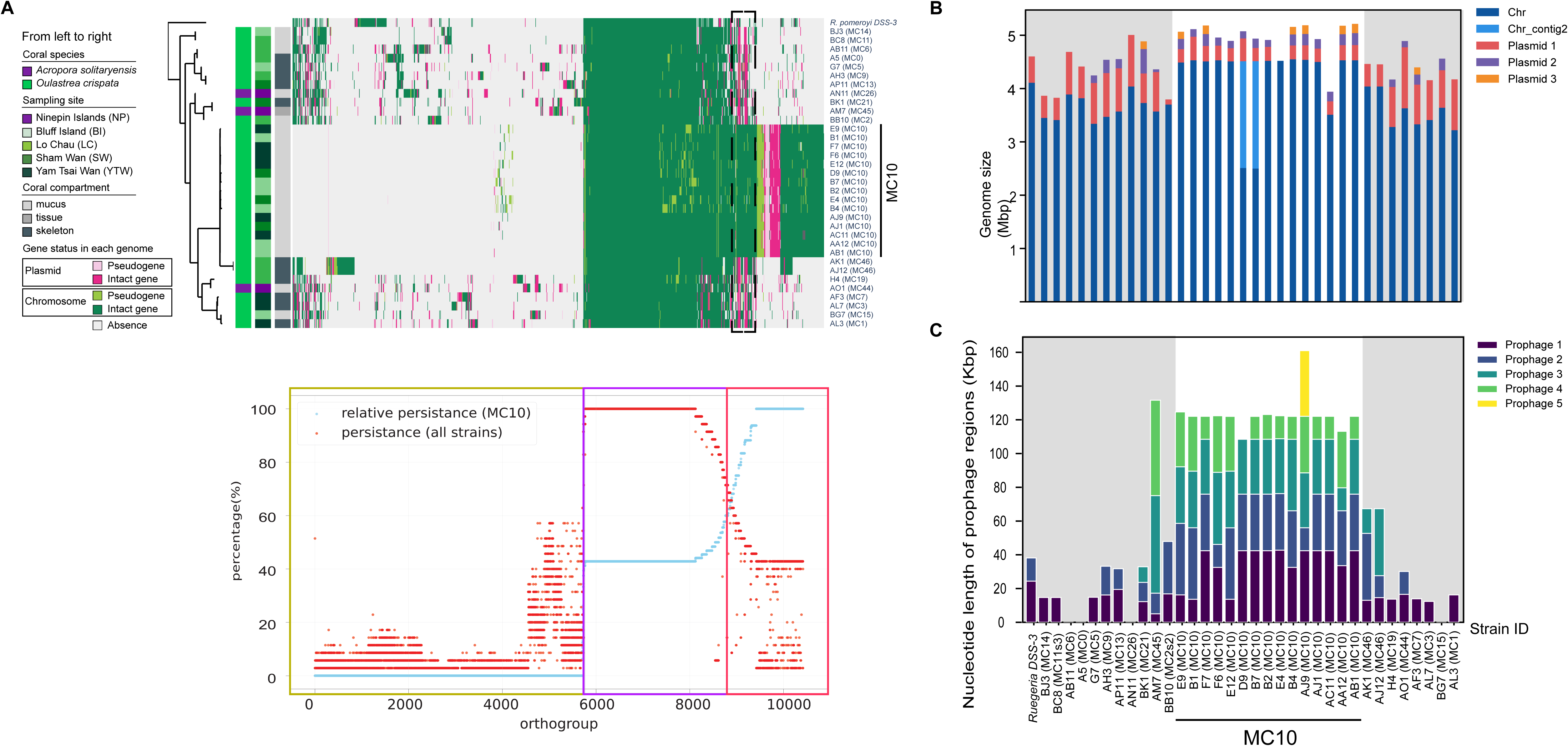
Overview of the 35 *Ruegeria* strains with closed genomes. (**A**) Upper: Presence/absence patterns of orthologous gene (OG) groups in chromosomes and plasmids, clustered by gene content variations. Three annotation columns next to the clustering dendrogram refer to *Ruegeria* isolates’ metadata: coral species, sampling site, and coral compartment, respectively. Gene status (pseudogene vs. intact gene) is color-coded by genomic location (chromosome/plasmid). The integrated chromosomal sequences in MC10 homologous to plasmids in non-MC10 *Ruegeria* are shown in a dashed frame. Lower: Scatter plot showing OG distribution across *Ruegeria* isolates. Each OG is represented by a single dot. Red dots (Global occurrence) show the percentage of all 35 isolates that contain a given OG, calculated as (number of genomes containing the OG) / (total 35 genomes). Blue dots (MC10-specific relative persistence) show a normalized measure of how concentrated an OG is within the MC10 population, calculated as (number of MC10 strains containing the OG)/(number of genomes containing the OG). Coloured outlines highlight key distributions: yellow for OGs rare in MC10 and sparsely distributed in non-MC10 *Ruegeria*, purple for OGs conserved in all populations, and red for OGs uniquely prevalent in MC10. (**B**) Stacked bar plot comparing chromosomal and plasmid sizes across the 35 strains based on Canu assemblies (∼4.5 Mb). MC10-D9 and MC10-B7 show nearly closed genomes (two contigs: the larger chromosome and a smaller contig labeled Chr_contig2), MC10-AC11 has a less complete assembly (∼3.51 Mb), while other MC10 members possess complete single-chromosome assemblies. (**C**) Stacked bar plot showing the distribution of integrated prophage lengths across strains, with each colored segment representing an individual prophage region (Prophages 1 to 5).

The genetic repertoire of *Ruegeria* MC10 was contextualized within the broader diversity of coral-associated *Ruegeria*. Clustering of the ∼10,400 gene families from the 35 closed genomes (4,119 ± 43 for each) revealed three broad genomic blocks: a core set of ∼3,200 gene families conserved across all populations, ∼5,700 gene families sparsely distributed in other *Ruegeria*, and ∼1,500 gene families distinctly prevalent in the MC10 population (Fig. 2A). Approximately 31% of MC10’s genomic regions are distinct and rarely found in other coral-associated *Ruegeria*.

The functional significance of MC10’s unique gene set was evident from significant enrichments in several KEGG modules, including cytochrome *bd* ubiquinol oxidase, deoxyribonucleotide biosynthesis, methionine degradation, pantothenate biosynthesis, and molybdenum cofactor biosynthesis (*P* < 0.05). In the reciprocal comparison, no KEGG modules were significantly enriched in non-MC10 *Ruegeria* relative to MC10 (Fig. S4).

The BinaryPGLMM analysis identified 116 genes with significantly differential abundance between MC10 and non-MC10 lineages, with 85 genes enriched in MC10 and 31 genes enriched in other *Ruegeria* (Dataset S3). MC10 showed enrichment for genes involved in host interaction and biofilm formation, including glycosyltransferases (GTs) from families GT51 (peptidoglycan glycosyltransferase, *P* < 0.001), GT0 (glycosyltransferases not yet assigned to a family, *P* < 0.001) GT2 (involved in sugar transfer to polysaccharides and lipids for biofilm matrix and capsule production, *P* > 0.05), and GT25 (lipopolysaccharide biosynthesis, *P* > 0.05) [21]. A cluster of genes for exopolysaccharide (EPS) production and secretion (*exoL*, *exoM*, *exoX*, *exoA*, *exoO*, *exoP*, *P* < 0.05; *exoU*, TC.PST, *P* = 0.06) and cell envelope modification (*gnu* for GlcNAc-P-P-Und epimerase, *P* < 0.001; *ompV* encoding an adhesin that facilitates bacterial attachment to host cells, *P* < 0.05) are enriched in MC10 (Fig. 3) [22–25]. MC10 uniquely harbors genes for specialized sugar metabolism, including *rhaM* (*P* < 0.05) for rhamnose metabolism and *fucD* (*P* < 0.01) for fucose metabolism, common components of bacterial EPS [26, 27]. MC10 exclusively possesses *icmK* encoding intracellular multiplication protein (*P* < 0.001), a component of the Dot/Icm Type IV secretion system known for translocating effector proteins into host cells to facilitate intracellular survival [28, 29].

**Fig. 3.**
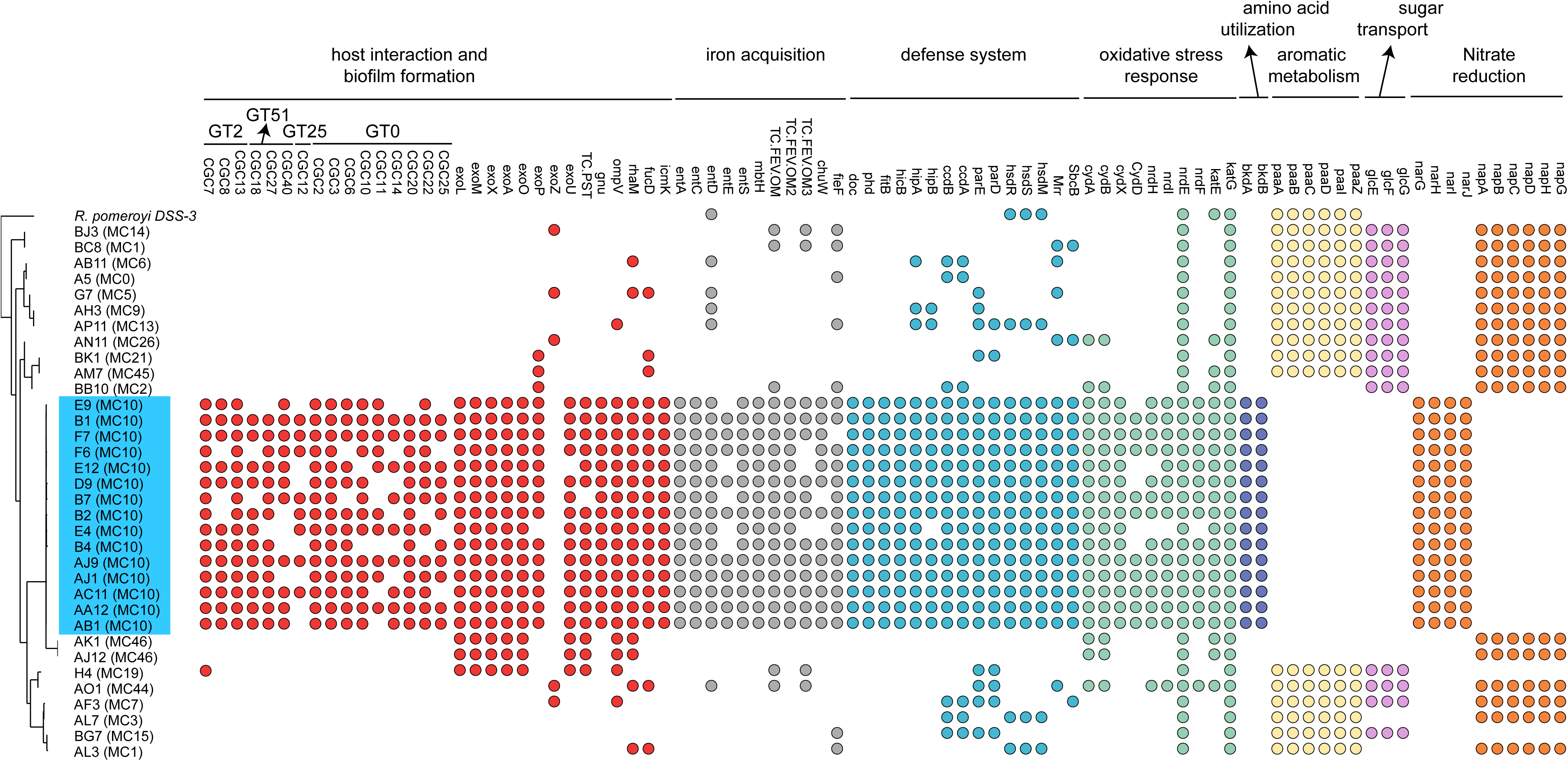
Functional gene distribution across the 35 closed *Ruegeria* genomes. Binary presence (filled) and absence (open) for each gene of interest are shown for all 35 *Ruegeria* genomes, with MC10 members highlighted in blue. Genes were selected based on functional relevance to host interaction and biofilm formation, iron acquisition, defense system, oxidative stress response, amino acid utilization, aromatic metabolism, sugar transport, and nitrate reduction.

For iron acquisition, MC10 carries a complete siderophore biosynthesis gene cluster (*entABCDEFS*) (Fig. 3) significantly enriched after phylogenetic control (*P* < 0.001 for most genes). This biosynthetic capability is complemented by outer membrane receptors, including TC.FEV.OM2 (KEGG ID: K16089, outer membrane receptor for ferrienterochelin and colicins, *P* < 0.001), TC.FEV.OM3 (K16087, hemoglobin/transferrin/lactoferrin receptor protein, *P* = 0.07), and TC.FEV.OM (K02014, iron complex outer membrane receptor protein, *P* >> 0.05). MC10 exclusively contains anaerobilin synthase (*chuW*, *P* < 0.01), an enzyme that enables oxygen-independent heme degradation, and is enriched in the ferrous-iron efflux pump *fieF* (*P* < 0.05).

MC10 showed significant enrichment for multiple defense systems. These include toxin-antitoxin modules *doc*-*phd* (*P* < 0.001), *fitB*-*hicB* (*P* < 0.001; although in MC10-B4 these two genes are not colocalized unlike typical toxin-antitoxin operons), *hipA*-*hipB* (*P* < 0.01 for *hipB*), *ccdB*-*ccdA* (*P* < 0.01), *parE*-*parD* (*P* = 0.05 to < 0.01) (Fig. 3). The functions of these toxins are diverse: *doc* inhibits protein synthesis and translation, *fitB* acts as an RNA-cleaving nuclease, *hipA* inactivates glutamyl-tRNA synthetase (GltX), *ccdB* targets DNA gyrase, and *parE* blocks DNA supercoiling [30]. The type I restriction enzyme system (*hsdRSM*, *P* < 0.01), restriction enzyme Mrr (*P* < 0.01), and exonuclease SbcB are also enriched in MC10.

Among oxidative stress-related genes, specific components showed significant enrichment. Within the cytochrome *bd* oxidase cluster, *cydX* was significantly enriched (*P* < 0.001), whereas *cydABD* were universally present. Within the Class Ib ribonucleotide reductase operon, *nrdFHI* were significantly enriched (*P* < 0.001), whereas *nrdE* was broadly present. Other oxidative stress genes present in MC10, including catalases *katE*, *katG*, and peroxidases, were not significantly enriched after phylogenetic control, suggesting their distribution reflects shared evolutionary history rather than lineage-specific selection.

Genes enriched in non-MC10 *Ruegeria* included the ring-1,2-phenylacetyl-CoA epoxidase operon (*paaABCDIZ*, *P* < 0.001), essential for aromatic compound metabolism via the phenylacetate pathway [31] and a glucose/mannose transport system (*glcEFG*, *P* < 0.001). For dissimilatory nitrate reduction, MC10 exclusively carries the NAR type (*narGHIJ*, *P* < 0.001), whereas non-MC10 strains exclusively carry the NAP type (*napABCDHG*, *P* < 0.001). MC10 was also significantly enriched in 2-oxoisovalerate dehydrogenase (*bkdAB*, *P* < 0.001), which enables utilization of branched-chain amino acids as carbon and energy sources.

### Phenotypic traits of *Ruegeria* MC10 are consistent with genomic predictions of a host-adapted symbiont

Genomic predictions translated into distinct realized cellular traits when comparing MC10-B4 against sympatric controls MC0-A5, MC15-BG7, and the free-living model *Ruegeria pomeroyi* DSS-3.

Transmission electron microscopy (TEM) revealed strain-specific differences in intracellular inclusions. Bright white storage granules, likely polyhydroxyalkanoates (PHAs), and light grey putative lipid droplets were prominent in DSS-3 and MC0-A5, reduced in MC10-B4, and absent in MC15-BG7 (Fig. 4A).

**Fig. 4.**
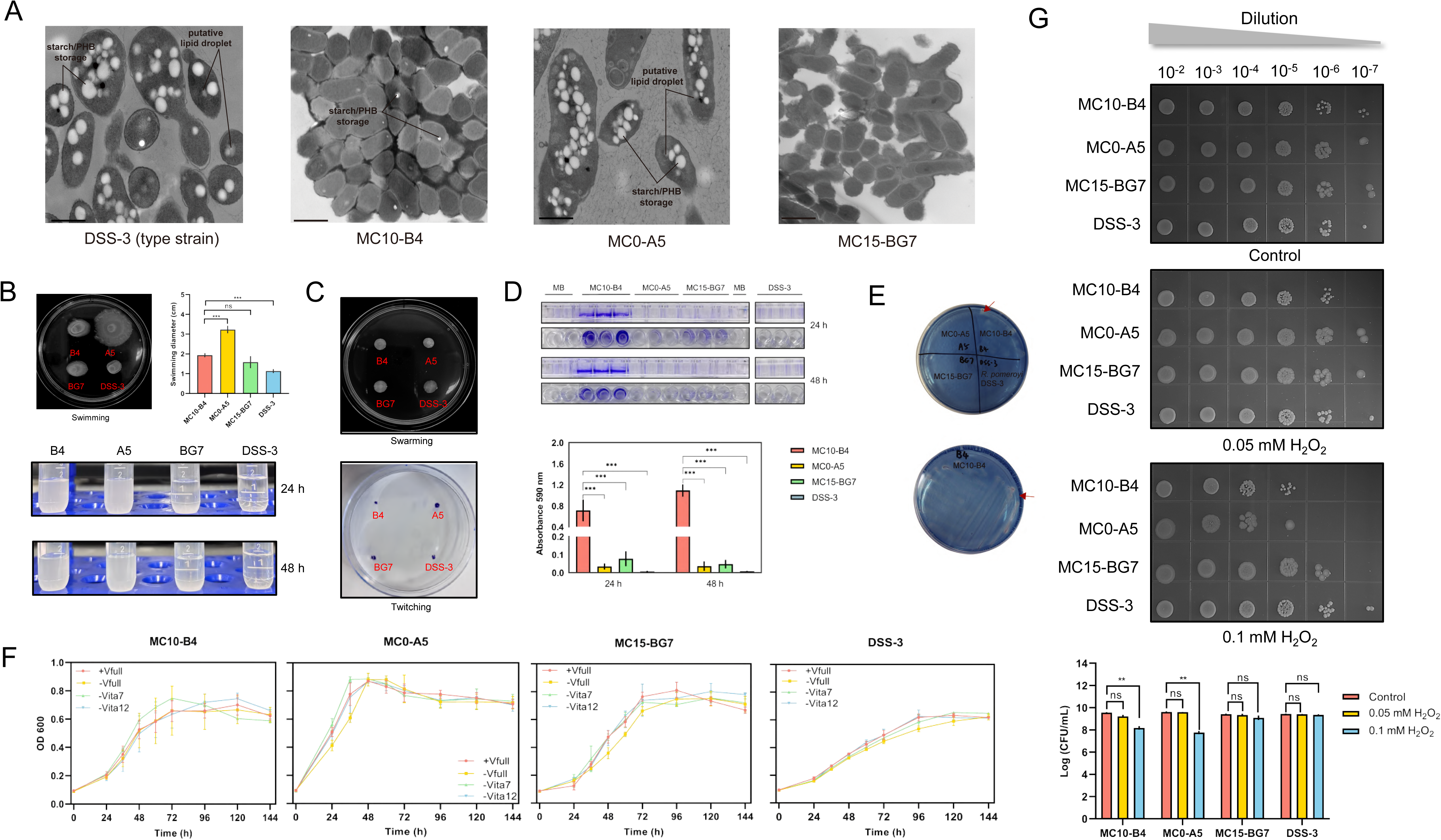
Morphological and functional characterization of four *Ruegeria* strains. (**A**) Transmission electron microscopy (TEM) images of *Ruegeria* type strain *R. pomeroyi* DSS-3, MC10-B4, MC0-A5, and MC15-BG7. Bright white granules correspond to intracellular storage inclusions, like starch and/or polyhydroxybutyrate (PHB), whereas light grey regions correspond to putative lipid droplets. Scale bars = 600 nm. (**B**) Bacterial swimming motility assay. (Top left panel) Representative images of swimming halos on 0.18% semi-solid marine broth 2216 agar plates after 6 days of incubation at 28 °C. Bacterial suspensions (5 μL) at logarithmic growth phase (OD_600_=0.5) were point-inoculated at the centre of each plate. (Top right panel) Quantification of swimming motility, presented as the diameter of the halo. Data are shown as mean ± SD from three independent experiments: ns, not significant, ***, *P* < 0.001. (Bottom panel) Sedimentation phenotypes of the four strains after 24 h and 48 h of incubation at room temperature without shaking. DSS-3 and MC15-BG7 show a clear sedimentation phenotype, whereas MC10-B4 and MC0-A5 did not settle under the same conditions. (**C**) Swarming and twitching motility assays. (Top panel) Bacterial suspensions (5 µL) of four strains at logarithmic growth phase (OD_600_=0.5) were spotted onto swarming plates (0.75% agar). (Bottom panel) For twitching motility, 1 µL of bacterial suspension was stab-inoculated into 1% agar plates. All plates were incubated at 28 °C for 10 days. To visualize twitching motility, the agar was carefully removed, and the adherent cells at the inoculation point were air-dried, fixed, and stained with 0.1% crystal violet. Images shown are from one experiment representative of three independent biological replicates. (**D**) Biofilm formation by four *Ruegeria* strains was quantified using crystal violet staining. After incubation in marine broth 2216 within a 96-well plate for 24 or 48 hours at 28 °C, the biofilm was stained with 0.1% crystal violet and subsequently decolorized with 0.2 ml decolorizing solution; absorbance was measured at OD_590_. All test strains were grown on the same plate. Data represent the mean of six biological replicates, with error bars indicating standard deviations. MB: Marine broth medium 2216. ***, *P* < 0.001. (**E**) Chrome azurol S (CAS) assay assessing siderophore production in the four *Ruegeria* strains. The red arrows point to the areas where the color changes from blue to yellow. (**F**) Growth of four *Ruegeria* strains assessed in marine basal medium (MBM) under different vitamin supplementation conditions. Four treatments were tested: (i) complete vitamin supplementation, (ii) no vitamins, (iii) absence of vitamin B_7_, and (iv) absence of vitamin B_12_. Cultures were inoculated at an initial OD_600_ of 0.01 and incubated at 28 °C with shaking. Growth was monitored by measuring OD_600_ over time. Data represent the mean of three replicates, and error bars indicate standard deviations. (**G**) Ten-fold serial dilutions of the four strains were spotted onto marine broth 2216 agar plates containing the indicated concentrations of H_2_O_2_ (0, 0.05, 0.1 mM). Plates were imaged after 3 days of incubation. The bottom panel quantifies the bacterial counts (Colony Forming Units, CFU) under each condition. Data are presented as the mean ± SD from three independent biological replicates. ns, not significant; **, *P* < 0.01.

Motility assays revealed that MC10-B4 displayed an intermediate phenotype, remaining in suspension after 48 hours (Fig. 4B), whereas MC0-A5 was highly motile with no sedimentation, and both DSS-3 and MC15-BG7 showed minimal motility with complete sedimentation within 24 hours (Fig. 4B). No swarming or twitching motility was observed for any strain after 10 days of incubation (Fig. 4C).

Biofilm formation assays showed that MC10-B4 developed substantially more robust biofilms than MC0-A5 and MC15-BG7 at both 24 and 48 hours, whereas DSS-3 formed no detectable biofilm under these conditions (Fig. 4D).

Iron chelation assays demonstrated a color change from blue to orange indicative of iron chelation only for MC10-B4; no color change was detected for MC0-A5, MC15-BG7, or DSS-3 (Fig. 4E).

Vitamin prototrophy assays in a defined marine basal medium showed that all four strains achieved comparable final biomass yields regardless of vitamin supplementation, including in media lacking B_12_ (cobalamin) or B_7_ (biotin), indicating prototrophy for these vitamins (Fig. 4F). Minor exceptions were observed for MC15-BG7, which showed a difference between complete vitamins and no B_12_ (one-way ANOVA; *P* < 0.05), and for DSS-3 showed a difference between no B_12_ and no B_7_ (*P* < 0.05); all other pairwise comparisons were non-significant.

Oxidative stress tolerance assays revealed that MC10-B4 showed reduced survival, with decreased colony formation at different H_2_O_2_ concentration compared to the other three strains (Fig. 4G, S5A). Catalase activity assays showed that MC15-BG7 vigorously produced bubbles; DSS-3 produced weak bubbles; MC10-B4 and MC0-A5 did not produce bubbles (Fig. S5B).

### Host metabolite exposure triggers a distinct motility[]to[]sessile transition in *Ruegeria* MC10

The functional response of *Ruegeria* MC10 to host-derived signals was elucidated through comparative proteomic profiling of strains MC10-B4, MC0-A5, and MC15-BG7 following exposure to macerated coral tissue extract. The analysis identified 6,267 expressed proteins across all conditions, with 5,528 high-confidence proteins retained after stringent filtering, clustered into 2,645 orthologous gene (OG) groups (Dataset S4).

The proteomic response to host metabolites was pronounced and comparable in scale across the three sympatric strains. Significant differential expression (FDR < 0.05, |log_2_FC| > 0.5) was observed in 583 of 2222 OGs (26%) in MC10-B4 (Dataset S5), 773 of 1980 OGs (39%) in MC0-A5 (Dataset S6), and 522 of 1793 OGs (29%) in MC15-BG7 (Dataset S7). The direction of regulation differed: MC0-A5 showed a strong predominance of upregulated proteins (590 up versus 183 down), whereas both MC10-B4 (191 up versus 392 down) and MC15-BG7 (182 up versus 340 down) were predominantly downregulated. Principal Coordinate Analysis (PCoA) revealed that bacterial strain identity was the primary driver of proteomic variation, explaining a significant proportion of the variance along PCoA1 (55.57%) and PCoA2 (37.75%) (Fig. 5A). The proteome of MC10-B4 was distinctly separated from those of MC0-A5 and MC15-BG7 along both axes, with the latter two strains separated from each other along PCoA2 but not PCoA1, underscoring the unique functional state of MC10-B4.

**Fig. 5.**
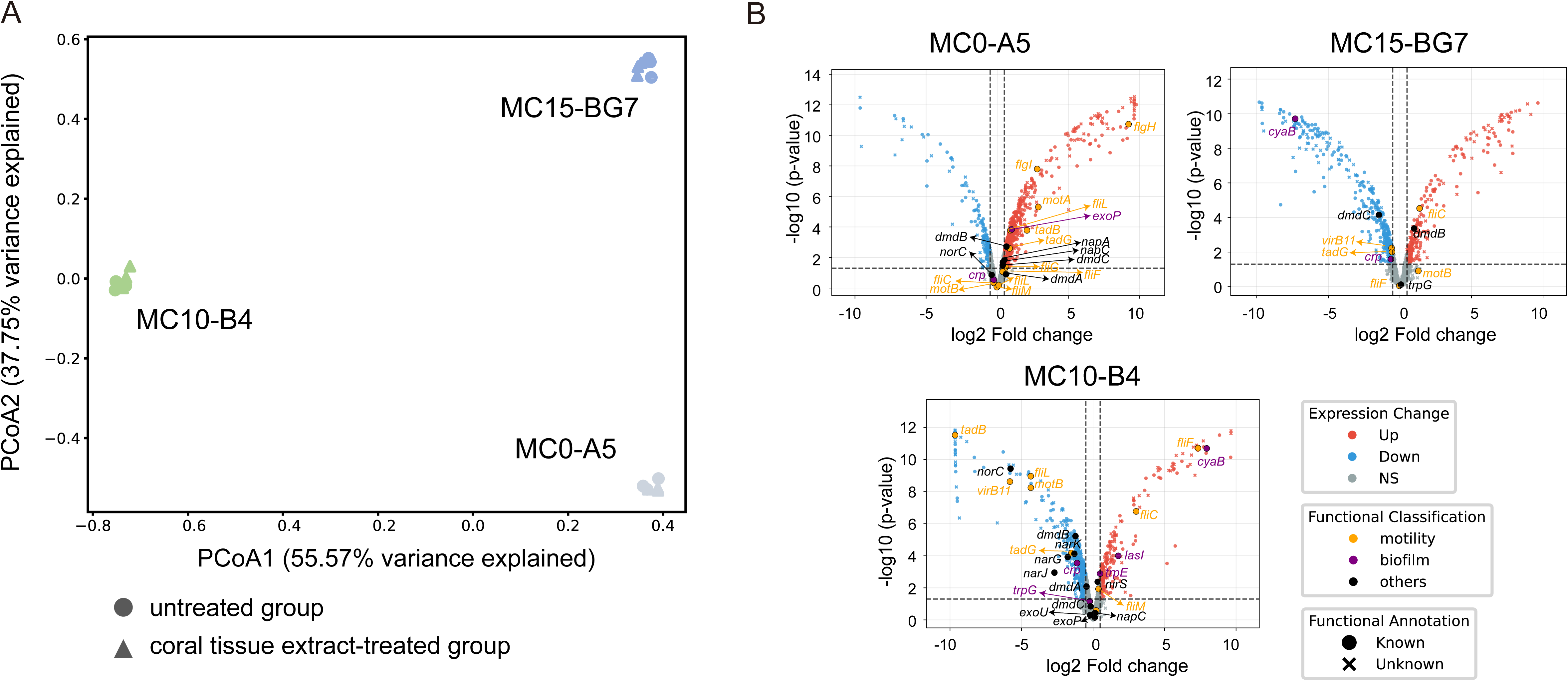
Proteomic analysis of coral tissue extracts effects on *Ruegeria* MC10-B4 and its ecological counterparts MC0-A5 and MC15-BG7. (**A**) Principal Coordinate Analysis (PCoA; Bray-Curtis dissimilarity) reveals distinct protein expression profiles, with MC10-B4 separating from MC0-A5 and MC15-BG7 along both PCoA axes (PCoA1 and PCoA2), while MC0-A5 and MC15-BG7 differ only along PCoA2. Treatment groups (coral tissue extract-treated vs. untreated) are indicated by shapes. (**B**) Volcano plot of differential gene expression. Each point represents a single gene, plotted by its log_2_ fold-change (log_2_FC) and -log_10_(P-value). Genes are coloured based on differential expression status: red for significantly upregulated (*P* < 0.05, log_2_FC > 0.5), blue for significantly downregulated (*P* < 0.05, log_2_FC < -0.5), and grey for non-significant features. Selected genes of interest are further classified by function: orange for motility, purple for biofilm formation, and dark for other functions. Point geometry denotes annotation status: circles (•) for genes with a functional annotation and crosses (×) for uncharacterized genes. The horizontal dash line indicates the P-value threshold (*P* = 0.05), and vertical dashed lines mark the fold-change thresholds (|log_2_FC| = 0.5).

Proteomic analysis revealed that exposure to coral tissue extract induced strain-specific changes in proteins associated with flagella-associated motility, with upregulation and downregulation defined by statistical significance (FDR < 0.05, |log_2_FC| > 0.5) (Fig. 5B, Dataset S8). In MC10-B4, flagellar proteins showed a mixed pattern of regulation. The flagellar M-ring protein FliF, which anchors the flagellum to the inner membrane, was upregulated, whereas the flagellar L-ring protein FlgH, which guides the flagellum through the outer membrane, was expressed but unchanged. The motor protein subunit MotB and the flagellar protein FliL were downregulated, whereas flagellin FliC was upregulated. In MC0-A5, a broad suite of flagellar and chemotaxis proteins was upregulated, including FlgH, the flagellar hook protein FlgE, the flagellar P-ring protein FlgI, FliL, the motor protein subunit MotA, and the chemotaxis protein CheY. FliF, MotB, and FliC were expressed but unchanged in MC0-A5. In MC15-BG7, FliC was upregulated, whereas FliF and MotB were expressed but unchanged. FlgE, FlgH, FlgI, FliL, MotA, and CheY were absent in MC15-BG7.

Pilus assembly and tight adherence (Tad) export apparatus proteins also showed strain-specific regulation (Fig. 5B, Dataset S8). In MC10-B4, the tight adherence protein B TadB was downregulated, whereas other pilus assembly proteins (CpaB, CpaC, CpaE, CpaF, TadC) were expressed but unchanged. MC0-A5 displayed a contrasting response, with upregulation of multiple pilus assembly proteins including CpaB, CpaC, CpaE, TadB, and TadC. CpaF was expressed but unchanged in MC0-A5. In MC15-BG7, all detectable pilus assembly proteins (CpaB, CpaC, CpaE, CpaF, TadC) were expressed but unchanged, whereas TadB was absent.

Biofilm-related proteins exhibited a mixed pattern of regulation upon host extract exposure (Fig. 5B, Dataset S8). The transcriptional regulator Crp, which mediates cyclic AMP (cAMP) signaling, was downregulated in MC10-B4 but expressed without change in MC0-A5 and MC15-BG7. Adenylate cyclase CyaB, which generates cAMP, was upregulated in MC10-B4 but downregulated in MC15-BG7. Anthranilate synthase TrpE, which supplies precursors for quorum sensing signals, was upregulated in MC10-B4, expressed but unchanged in MC0-A5, and downregulated in MC15-BG7. Serine O-acetyltransferase CysE, which can influence biofilm formation through cysteine metabolism, was upregulated in MC0-A5 but expressed without change in MC10-B4 and MC15-BG7. Phenylacetate-CoA ligase PaaK, part of a degradation pathway linked to biofilm formation and oxidative stress resistance, was upregulated in MC15-BG7 but expressed without change in MC10-B4 and MC0-A5. The polysaccharide biosynthesis transport protein ExoP, critical for exopolysaccharide polymerization and export during biofilm matrix assembly, was present only in MC10-B4 and MC0-A5; it was upregulated in MC0-A5 but expressed without change in MC10-B4.

Beyond motility and biofilm, coral tissue extract induced widespread metabolic reprogramming revealing distinct strategies between strains. Analysis of denitrification pathways, informed by the genomic presence of complete denitrification machinery in MC10-B4 and MC0-A5 but not in MC15-BG7, showed divergent expression patterns. In MC10-B4, the membrane-bound nitrate reductase gene *narG* was downregulated, *narH* was expressed but unchanged, and *narI* was not expressed. The nitrite reductase *nirS* was upregulated, the nitric oxide reductase *norC* was downregulated, and *norB* was not expressed. Nitrous oxide reductase *nosZ* was expressed but unchanged. In MC0-A5, which carries the periplasmic nitrate reductase pathway, *napA* was upregulated and *napB* was expressed but unchanged; *nirS* was expressed but unchanged; *norC* was expressed but unchanged and *norB* was not expressed; *nosZ* was upregulated. MC15-BG7, which genomically lacks denitrification machinery, showed no detectable expression of these pathways.

The xanthine dehydrogenase (*xdhABC*) and uric acid degradation pathway exhibited strain-specific regulation. MC10-B4 downregulated *xdhBC* and the 5-hydroxyisourate hydrolase gene (K07127), which converts uric acid to allantoin, while leaving other pathway genes unchanged. MC0-A5 upregulated *xdhC* and the ureidoglycine aminohydrolase gene (K14977), downregulated the allantoicase gene (K16842), and left other genes unchanged. MC15-BG7 downregulated K14977 with other genes unchanged.

Analysis of DMSP catabolism, for which all three strains possess the demethylation pathway (*dmdABC*) and acrylate detoxification gene *acuI* but lack cleavage pathways (e.g., *ddd* genes), revealed differential expression. MC15-BG7 additionally possesses *dmdD*, the final enzyme in the demethylation pathway. MC10-B4 downregulated *dmdAB*, while *dmdC* remained unchanged. MC0-A5 upregulated *dmdBC* and *acuI*. In MC15-BG7, the key gene *dmdA* was not expressed.

Nitrogen acquisition and regulatory pathways showed contrasting patterns between strains. MC10-B4 downregulated several nitrogen assimilation genes, including the proline aminopeptidase *pepP*, components of the general L-amino acid transport system (*aapQMP*), the branched-chain amino acid transport system (*livG*), glutamine synthetase *glnA*, glutamate synthase *gltB*, the nitrogen regulation two-component system sensor *ntrY*, and the spermidine/putrescine transport system *potB*. Other nitrogen-related genes were unchanged or unexpressed. MC15-BG7 similarly downregulated *pepP*, alanine aminopeptidase *pepN*, methionine aminopeptidase *pepM*, amino acid transport components (*aapQP*, *livG*), *glnA*, *potA*, and the

polyamine transport gene *potG*, with other genes unchanged or unexpressed. In contrast, MC0-A5 upregulated multiple nitrogen acquisition genes, including leucine aminopeptidase *pepL*, *pepP*, *pepN*, amino acid transport components (*aapQM*, *livK*), the P-II nitrogen regulator *glnB*, the two-component system regulators *ntrB* and *ntrX*, and *potG*, while downregulating *livF*, *glnA*, and *ntrY*, with other genes unchanged or unexpressed.

## Discussion

### *Ruegeria* MC10’s enhanced probiotic efficacy validates evolution-guided selection framework

The evolution-guided framework proposed in the companion study predicted and validated that candidates selected for genomic signatures of emerging host dependency exhibit long-term host association [4]. Whether this persistent colonization translates to discernable host benefits, however, requires direct testing. To address this, we compared thermal tolerance changes following incubation of three sympatric, coral-derived *Ruegeria* strains isolated from the same coral colony and mucus compartment. This design controls for host genotype and micro-environment, as probiotic engraftment is known to be strain-specific and shaped by host factors [32]. In standardized heat-stress assays using the model cnidarian Aiptasia H2, *Ruegeria* MC10-B4 inoculation significantly increased host thermal tolerance compared to a seawater placebo, with a 1.0 °C higher ED50 immediately post-stress and a 0.6 °C higher ED50 after recovery (Fig. 1). MC0-A5 provided intermediate protection, elevating ED50 by 0.6 °C at T7 but showing no significant benefit after recovery. MC15-BG7 conferred minimal benefit, with no detectable effect at T18. Only MC10-B4 significantly improved thermal tolerance after the recovery phase, indicating that its protective effect retains functional capacity of algal endosymbionts rather than merely delaying acute stress damage. These results confirm that the genomic signatures of emerging host dependency identified in MC10 are associated not only with enhanced, long-term colonization [4] but also with discernable host benefits compared to sympatric strains.

### A coordinated multi-omics profile underpins MC10’s host-adapted lifestyle

*Ruegeria* MC10 possesses a distinct genome architecture consistent with an early-stage evolutionary transition toward host dependency [4]. The population was initially identified based on two canonical signatures: insertion sequence expansion and pseudogene proliferation, proposed to be hallmarks of host transitioning [5]. Comparative genomics reveal additional features that further support this trajectory. MC10 exhibits a chromosome enriched with integrated plasmid-derived sequences and a higher burden of intact prophages, features absent from its sister population MC46 and from other coral-associated *Ruegeria*. Recent models of insect endosymbionts (e.g., *Sodalis*, *Arsenophonus*) suggest that the initial shift to endosymbiosis is characterized by genome expansion driven by mobile genetic element proliferation, which can provide novel adaptive traits [33]. Consistent with this model, *Ruegeria* MC10’s genomes show hallmarks of this genome expansion phase. Nevertheless, MC10 presents a notable nuance: it has acquired and maintained a diverse arsenal of defense systems, including restriction-modification modules and multiple toxin-antitoxin families, which are significantly enriched after phylogenetic control. This observation suggests that the path to host dependency does not universally require comprehensive disarmament. MC10 may instead represent a state where certain mobile elements are harnessed while core defensive capabilities are retained for stability in a still-competitive host environment. Thus, whereas MC10’s genome expansion firmly places it in the early, plasticity-dominated phase of symbiotic transition, its uniquely acquired defense repertoire highlights an alternative trajectory within this model.

Integrating genomic predictions with phenotypic validation revealed three distinct patterns of concordance that inform how genomic signatures should be interpreted. Strong concordance was observed for siderophore production: genomic analysis revealed that MC10 possesses a complete siderophore biosynthesis cluster significantly enriched after phylogenetic control, and this prediction was robustly confirmed by phenotypic assays, with MC10-B4 being the only strain that produced detectable iron chelators in the chrome azurol S assay (Fig. 4E). This concordance validates siderophore-mediated iron acquisition as a core, realized trait contributing to persistence in the iron-limited host environment. Discordance was observed for vitamin auxotrophy. Genomic analysis suggested potential auxotrophy for vitamins B_7_and B_12_ due to pseudogenization, yet growth assays showed that MC10-B4, like all comparison strains, achieved comparable final biomass yields regardless of vitamin supplementation (Fig. 4F). Proteomic analysis added further nuance: upon host metabolite exposure, MC10-B4 expressed the full suite of B_12_ biosynthetic proteins despite its pseudogenes, whereas control strains with genomically intact pathways did not express several corresponding proteins. This finding suggests that host cues can trigger expression even from compromised loci. A third pattern, context-dependent discordance, emerged for oxidative stress defense. Genomically, MC10 carries an extensive arsenal of oxidative stress genes, including specific components of cytochrome *bd* oxidase (*cydX*) and the Class Ib ribonucleotide reductase operon (*nrdFHI*) that are significantly enriched after phylogenetic control (Fig. 3). MC10-B4 possesses both *katE* and *katG*, whereas sympatric control strains lack *katE* entirely and possess only *katG*. Phenotypically, MC10-B4 was distinctly sensitive to H_2_O_2_ and lacked detectable catalase activity under standard laboratory conditions (Fig. 4G, S5). Examination of proteomic data revealed that this sensitivity is not due to a simple lack of expression: all catalase genes present in a given strain were constitutively expressed under both control and host-cue exposure conditions, with no differential regulation observed. This context-dependent discordance carries an important implication: even constitutive expression of a full genetic arsenal *in vitro* does not translate into phenotypic trait manifestation. Screening strategies that rely solely on laboratory performance against H_2_O_2_ would have eliminated MC10-B4 from consideration, yet its oxidative stress repertoire likely contributes to successful long-term residency in the host environment.

Proteomic profiling revealed that the three sympatric strains adopted fundamentally different strategies in response to host tissue extract. MC10-B4 responded with a coordinated motility-to-sessility shift, downregulating multiple flagellar components (*fliL*, *motB*) and the tight adherence protein (*tadB*) while upregulating flagellin FliC and select biofilm regulators (*cyaB*, *trpE*) (Fig. 5B). FliC has dual functions in motility and initial surface attachment; its upregulation alongside downregulation of other motility machinery suggests a transition toward a sessile state [34, 35]. TadB downregulation parallels observations in *Bradyrhizobium diazoefficiens*, where Tad pili deletion increased adhesion and impaired motility [36]. Downregulation of cAMP receptor protein (*crp*) in MC10-B4 matches biofilm-promoting strategies in *Serratia marcescens* and *Vibrio cholerae* where CRP suppresses biofilm formation [37], whereas upregulation of adenylate cyclase (*cyaB*) promotes capsule production and biofilm matrix development [38]. By contrast, MC0-A5 mounted broad upregulation of motility, adhesion, and nutrient acquisition systems, including flagellar components, chemotaxis proteins, pilus assembly proteins, and serine O-acetyltransferase CysE; in *E. coli*, cysE mutants exhibit accelerated biofilm formation, suggesting CysE upregulation may suppress biofilm development [39], treating host metabolites as a nutritional resource rather than a colonization cue. This interpretive divergence extended across multiple metabolic pathways. In DMSP metabolism, MC10-B4 downregulated the demethylation pathway (*dmdAB*). DMSP is an abundant sulfur compound produced by coral host and symbionts [40, 41] with established antioxidant properties [42]; by minimizing its catabolism, MC10-B4 may preserve this molecule for host-level oxidative stress protection rather than harvesting it as a carbon and sulfur source. In purine degradation, MC10-B4 suppressed xanthine dehydrogenase subunits (*xdhBC*) and the uric acid conversion enzyme 5-hydroxyisourate hydrolase. Xanthine dehydrogenase catalyzes xanthine oxidation to uric acid, a reaction that can generate superoxide and hydrogen peroxide as byproducts, whereas uric acid and its downstream product allantoin possess ROS-scavenging properties [43–45]. Downregulation of these genes therefore reduces both endogenous ROS generation and catabolism of antioxidant compounds, a pattern consistent with prioritizing ROS mitigation. In nitrogen acquisition, MC10-B4 downregulated multiple aminopeptidases, amino acid transporters, and regulatory systems (*glnA*, *ntrY*) for several nitrogen acquisition pathways, whereas MC0-A5 upregulated these same pathways for active nitrogen scavenging. Denitrification expression further distinguished the strains. MC10-B4 carries the membrane-bound NAR-type nitrate reductase and upregulated nitrite reductase *nirS* while suppressing nitrate reductase *narG* and nitric oxide reductase *norC*. This expression pattern favors nitric oxide (NO) production over complete denitrification to dinitrogen. NO is increasingly recognized in beneficial symbioses as a signaling molecule that modulates host-microbe interactions [46]. MC0-A5, by contrast, upregulated nitrate reductase (*napA*) and nitrous oxide reductase (*nosZ*), a pattern consistent with complete denitrification for energy conservation. This coordinated program, which prioritizes colonization signaling, ROS mitigation, and metabolite preservation over nutrient harvesting, was unique to MC10-B4 and provides a mechanistic explanation for its superior persistence and host benefit.

For a symbiont to achieve long-term residency, it must navigate host-derived stress responses and competition from other microbes. Isolation from mucus does not definitively indicate a bacterium’s primary colonization site, as shed Symbiodiniaceae cells and their tightly associated bacteria can be present in the mucus layer, and mucus itself may serve as an initial contact site before tissue entry. Our *ex hospite* analyses therefore provide independent evidence that MC10 employs niche specialization complemented by genomic plasticity. Phenotypically, MC10-B4’s sensitivity to H_2_O_2_ indicates avoidance of high-ROS microenvironments such as the direct vicinity of Symbiodiniaceae, a finding consistent with FISH imaging showing that MC10 cells do not consistently co-localize with algal symbionts [4]. Genomically, this interpretation is supported by features such as the complete Class Ib ribonucleotide reductase operon for DNA synthesis under oxidative stress, anaerobilin synthase for iron acquisition in oxygen-limited environments [47], and the NAR-type nitrate reductase suggesting adaptation to specific oxygen regimes [48, 49]. The arsenal of defense systems uniquely acquired by MC10, including toxin-antitoxin modules and restriction-modification systems, equips MC10 with mechanisms to maintain genomic integrity against mobile genetic elements and phage attacks, potentially enhancing its stability as a probiotic inoculant [10, 50]. This combination of niche selection, metabolic integration, and genomic defense mechanisms may mitigate the inherent costs of MC10’s metabolic dependencies, facilitating sustained persistence.

### Rethinking probiotic selection: what MC10 teaches us about host adaptation

The functional profile of *Ruegeria* MC10 stands in contrast to traits typically selected for in conventional probiotic screens. Traditional approaches often prioritize bacterial performance *in vitro*, operating on the assumption that traits like high antioxidant activity or rapid nutrient cycling will function similarly within the host environment. Resistance to oxidative stress is a commonly selected trait [3, 51], predicated on the assumption that probiotics mitigate host bleaching by scavenging ROS. Counter to this expectation, MC10-B4 is uniquely sensitive to H_2_O_2_, a trait consistent with avoidance of the high-ROS symbiosome niche, noting that baseline ROS production is an inherent feature of coral holobionts [52, 53]. The presence of genes such as the DMSP demethylase *dmdA*, commonly used as a proxy for DMSP demethylation capacity, has been interpreted as indicative of beneficial function [3]. However, MC10-B4 downregulated this pathway upon host exposure. This suggests that such proxies can be misleading, and that traits which would exclude MC10 from conventional screens may instead underpin its successful host adaptation. The proteomic data further demonstrate that MC10’s colonization success derives not from superior nutrient harvesting but from a coordinated program that prioritizes host attachment, signaling, and ROS mitigation over nutrient acquisition. Thereby, this validation supports an evolution-guided approach that selects for innate colonization potential and host integration capacity over pre-defined *in vitro* functionalities.

The principle of selecting for evolutionary predisposition to host association offers a generalizable strategy for microbiome engineering. Medicine, aquaculture, agriculture, and conservation share the challenge of transient probiotic colonization. This study demonstrates that a candidate identified through genomic signatures of emerging host dependency possesses a functionally coordinated suite of traits, including siderophore-mediated iron acquisition, constitutive biofilm formation, and a host-responsive program prioritizing colonization over nutrient harvesting, which together enable stable persistence and discernable host benefits. By first identifying lineages with genomic signatures of long-term host association in our companion study and then characterizing their functional traits and empirically testing efficacy in this work, the framework validated here provides a scalable pathway toward durable, predictably effective probiotic interventions across diverse host systems.

## Acknowledgements

We would like to thank Xiaojun Wang and Wudi Gai for suggestions on bioinformatics, and Yumei Chen, Zhihao Xian, Jeremy Ho, and Kai Yan Lai for their support in phenotypic and morphological experiments.

## Data availability

Raw genome sequencing data are available on NCBI under several BioProjects with the private access link for reviewers. The raw reads of 409 and assembled genomes of 419 *Ruegeria* isolates (10 genomes missing raw data) sequenced on the second-generation platform (DNBseq) and 34 *Ruegeria* isolates sequenced on the Nanopore platform are available under NCBI BioProject accessions PRJNA1264799 (https://dataview.ncbi.nlm.nih.gov/object/PRJNA1264799?reviewer=mp7rulknapinfb95c9733nf72r) and PRJNA1275854 (https://dataview.ncbi.nlm.nih.gov/object/PRJNA1275854?reviewer=g4mffnb8b965tq6b2662cpt6i8), respectively. The mass spectrometry output files have been deposited in the ProteomeXchange Consortium via the PRIDE partner repository with the dataset identifier PXD069552 (https://www.ebi.ac.uk/pride/archive/projects/PXD069552).

## Funding

This study is supported by Hong Kong Research Grants Council (RGC) General Research Fund (project #: 14114724), along with a Direct Grant from CUHK Faculty of Science (project #: 4053691), granted to H.L. C.P. was supported by an ANR Junior Professor Grant ‘A connected underwater world’ ANR-22-CPJ2-0113-01 awarded by the French National Research Agency (ANR). C.R.V. acknowledges funding from the German Research Foundation (DFG) project no. 458901010 (https://gepris.dfg.de/gepris/projekt/458901010?language=en).

## Notes

### Competing Interest Statement

The authors have declared no competing interest.

### Summary of Updates

Figure 1 added. Dataset S8 added.

