## Supplementary figures for "From genomic decay to functional advantage: Traits of a persistent, thermally beneficial coral probiotic"

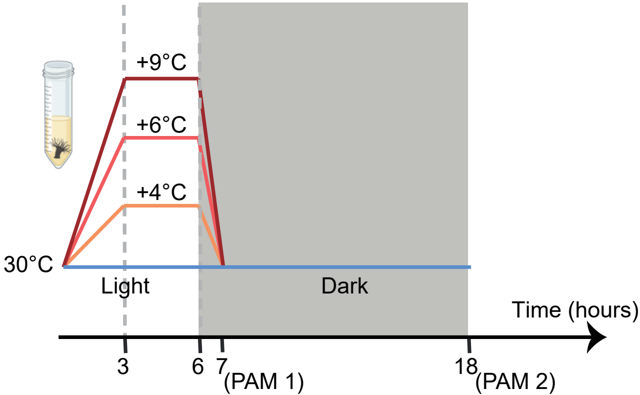


**Fig. S1.** Temperature profiling of the CBASS.

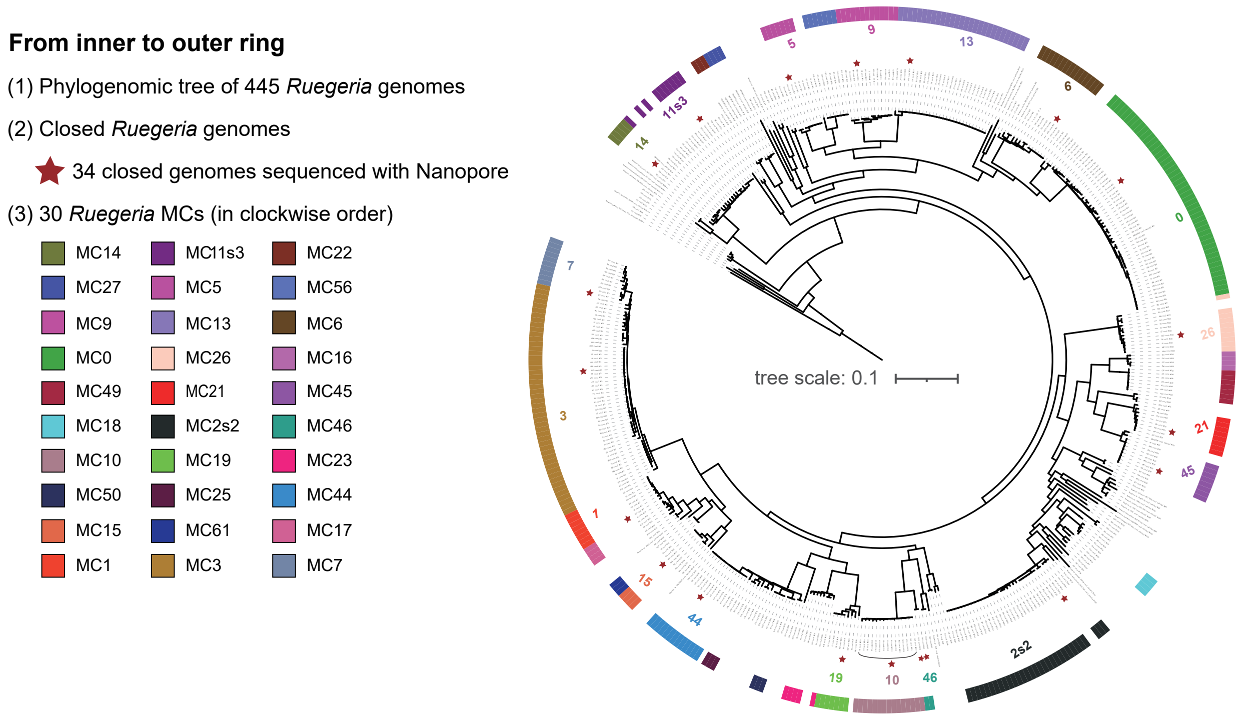


**Fig. S2**. **Rooted maximum-likelihood phylogenomic tree of 445 *Ruegeria* strains.** From inner to outer ring: (1) The phylogenomic tree comprising 419 newly isolated coral-associated *Ruegeria* strains and 26 publicly available genomes (NGS genomes). This tree was constructed with IQ-TREE based on concatenated single-copy orthologous genes at the amino acid level. (2) 34 closed genomes sequenced with Nanopore were labelled with stars in the tree. (3) PopCOGenT was used to delineate the *Ruegeria* populations (i.e., main clusters or MCs). 29 *Ruegeria* populations which consists of at least three isolates as defined by PopCOGenT, along with MC46, the sister group of MC10, comprising two isolates.

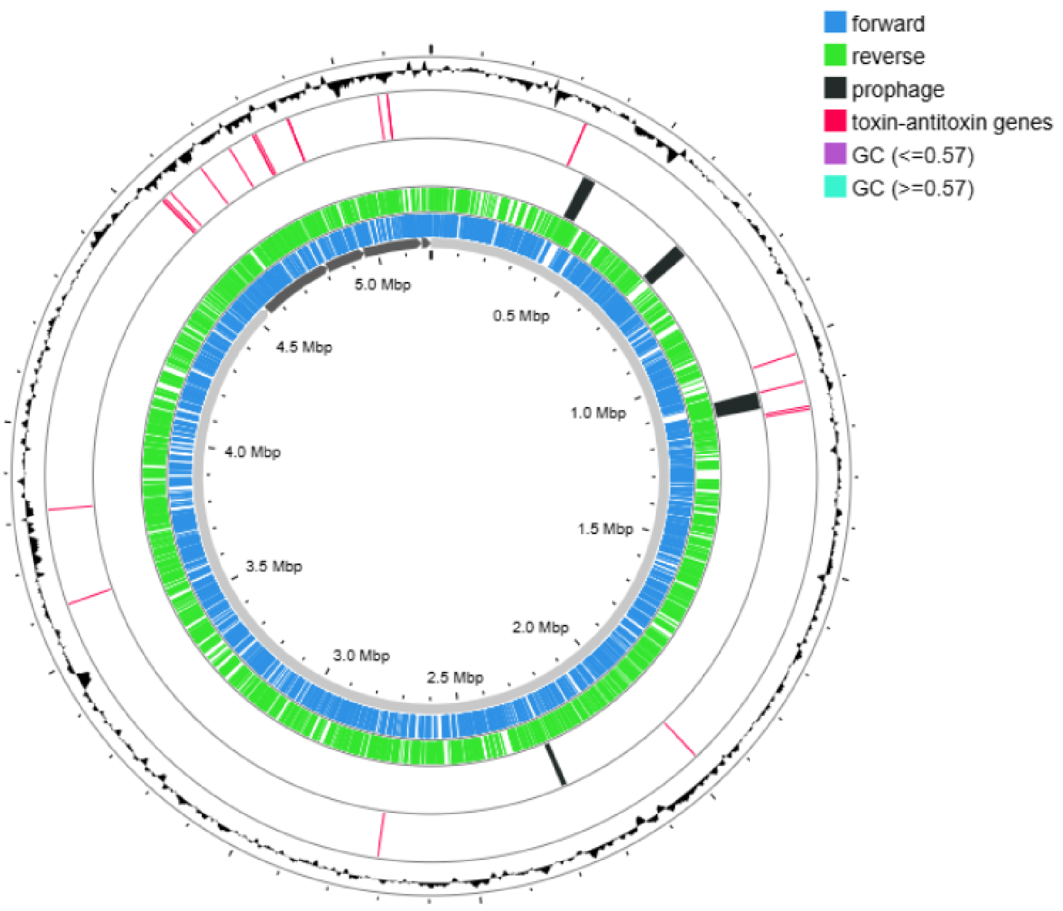


**Fig. S3. Genome maps of *Ruegeria* MC10-B4.** From inner to outer ring: (1) chromosomes and plasmids; (2) annotation of predicted genes on the forward strand; (3) annotation of predicted genes on the reverse strand; (4) putative prophages, (5) toxin-antitoxin genes; (6) GC skew (outer: GC ≥ 0.57; inner: GC < 0.57).

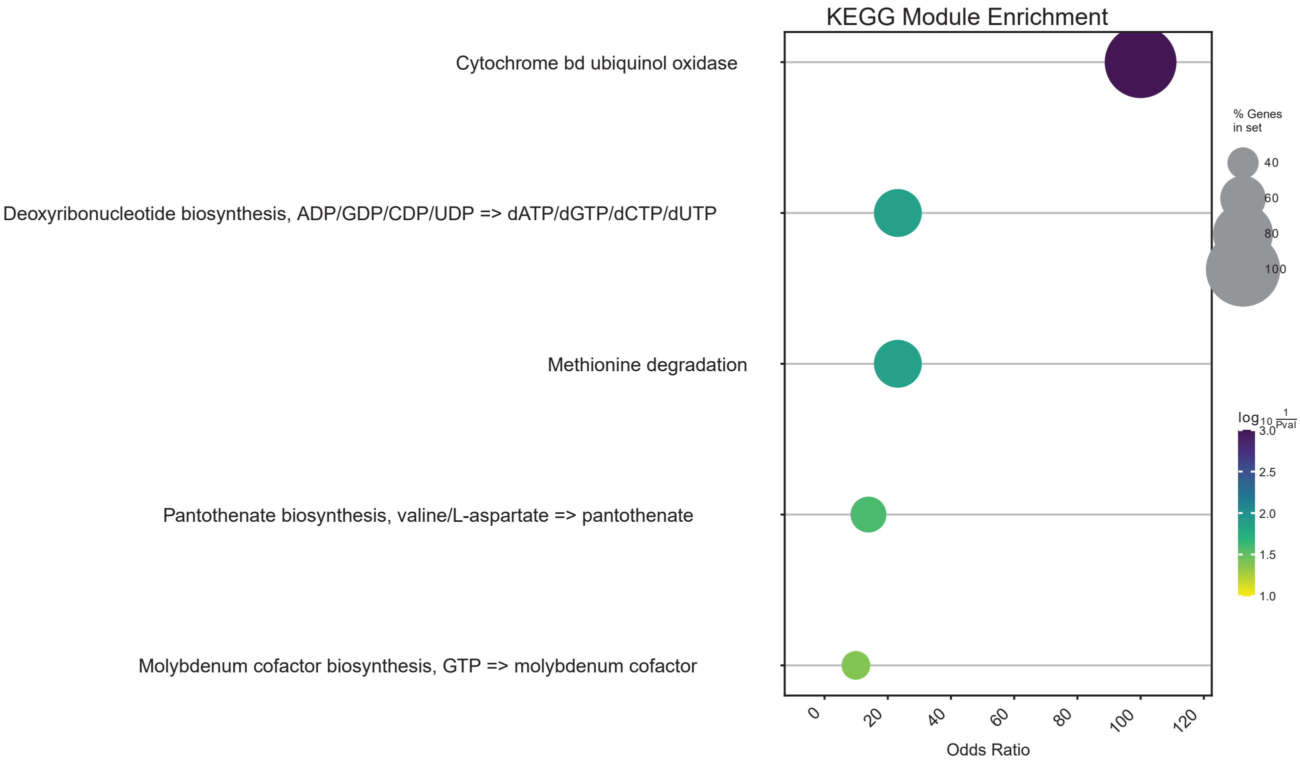


**Fig S4. KEGG module enrichment analysis of MC10-specific genes.** The bubble plot illustrates functional modules significantly enriched in the MC10 members (n=15) compared to the other *Ruegeria* populations (n=20). Y-axis shows KEGG Module Descriptions; X-axis represents the Odds Ratio, indicating the strength of functional enrichment. The dot size is proportional to the number of gene hits, reflecting functional dosage. The color indicates the Adjusted P-value derived from GSEApy (hypergeometric test). All displayed moduless satisfy the dual criteria of GSEApy P-value < 0.05 and PGLS P-value < 0.05.

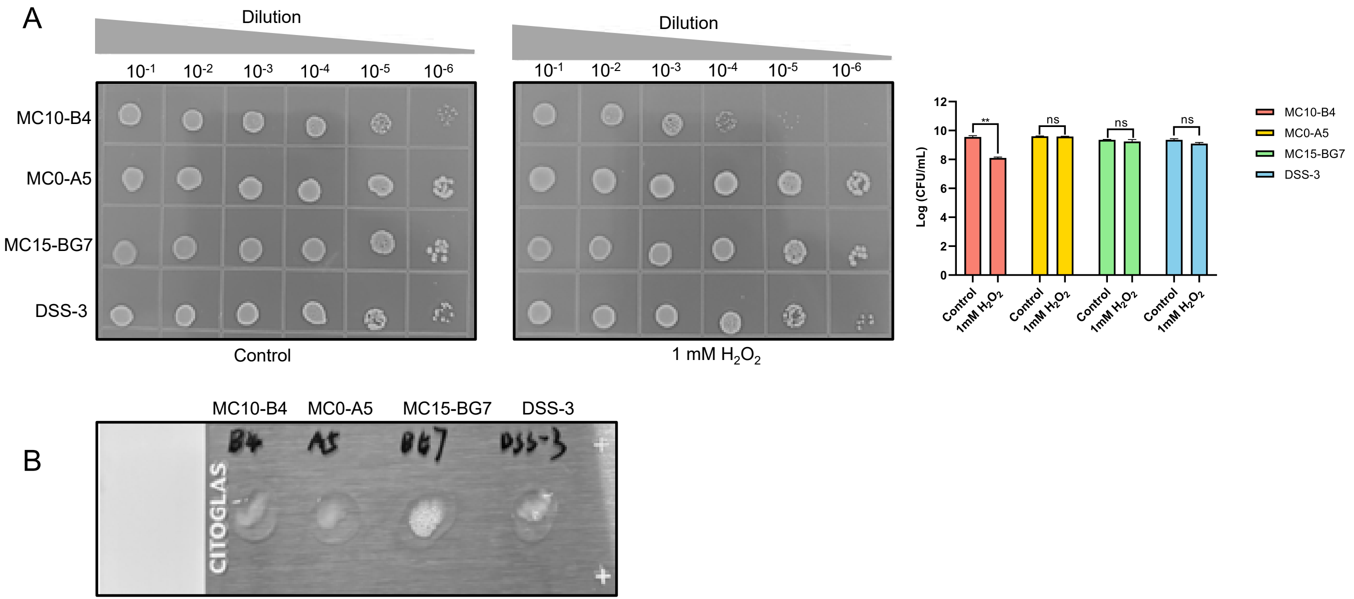


**Fig. S5 $H_2O_2$ sensitivity and catalase activity in *Ruegeria* strains.** (**A**) The ability of the indicated strains to survive in the presence and absence of 1 mM $H_2O_2$. Exponential-phase cultures were harvested, adjusted to an $OD_{600}$ of 0.5, and then incubated for 1 hour at 28 °C with or without (control) 1 mM $H_2O_2$. After incubation, cells were serially diluted, spread on marine broth 2216 agar , and incubated at 28 °C for 3 days. The representative plate images are shown. The right panel quantifies the bacterial counts (Colony Forming Units, CFU) from three independent experiments. Data presented as mean ± SD. **, $P < 0.01$; ns, no significant. (**B**) Catalase activity was directly visualized using a slide catalase test. Bacterial colonies were transferred to a glass slide and mixed with 10 µL 3% $H_2O_2$. The immediate formation of oxygen bubbles was observed and photographed, indicating the presence of catalase enzyme.
