## Supplementary information for "From genomic decay to functional advantage: Traits of a persistent, thermally beneficial coral probiotic"

**This PDF file includes:**

**Supplementary methods**

**Supplementary references**

**Figures S1 to S5**

**Supplementary methods**

***Ruegeria* isolation, genome sequencing, and analysis**

To generate data for analysis, we followed a streamlined workflow involving bacterial isolation from coral samples, genomic DNA extraction, and genome sequencing, assembly and annotation. This method allowed us to characterize the genomic features of these coral-associated *Ruegeria* strains and their potential functional roles. For a detailed description of the protocols and bioinformatic workflows, please refer to our previous paper [1], which presents the full methodological framework.

**Coral Bleaching Automated Stress System (CBASS) assays**

CBASS experiments were conducted on the anemone model Aiptasia H2 following inoculation with *Ruegeria* MC10-B4, MC0-A5, or MC15-BG7 (10^6^ CFU/mL in 0.22 μm-filtered seawater (FSW), inoculated 24 hours before CBASS initiation and kept throughout the 18 h of CBASS run) [2]. Given the capacity of the CBASS system, we conducted two rounds of CBASS runs using the anemones maintained in the same condition (salinity: 35 ppt; temperature: 25 °C; light condition: 12 h light: 12 h dark diurnal cycle, 70-80 μmol m^-2^ s^-1^ PAR) for 3 months. Animals were fed once a week with freshly hatched *Artemia* nauplii (Marine Nutrition, Inc.). Tanks were cleaned and supplied with fresh artificial seawater the next day. The first CBASS tested the efficacy of MC10-B4, MC0-A5, and the second tested the efficacy of MC15-BG7 on affecting the ED50 thermal threshold tolerances of Aiptasia H2. A placebo FSW anemone cohort was included for each run as a baseline thermal tolerance threshold (ED50) for the Aiptasia H2 without bacterial inoculation. For each treatment in the first CBASS run, 20 anemones were distributed across four 10-L aquariums (30 °C control, 34 °C medium stress, 36 °C high stress, 39 °C extreme stress; 5 replicates each) with each anemone maintained in a separate 50 mL centrifuge tube [2, 3]. For each treatment in the second CBASS run, 16 anemones were distributed across four 10-L aquariums (30 °C control, 34 °C medium stress, 36 °C high stress, 39 °C extreme stress; 4 replicates each), given limited anemone availability. Each aquarium contained a full-spectrum LED lighting (Galaxyhydro, China), a programmable thermostat (Inkbird ITC-310 T-B, China), an iceProbe thermoelectric cooler (Nova Tec, China), and a 200 W Titanium aquarium heater (Schego, China). Control tanks were maintained at 30 °C throughout the 18-hour experiment; stress profiles ramped to target temperatures over 3 hours, held at the target temperatures for 3 hours, returned to 30°C within 1 hour, then maintained for 11 hours (see Figure S1 for the thermal cycling profiles). Experiments ran overnight (1 p.m.–7 a.m., University of Konstanz, Germany) to avoid daylight interference, with temperatures logged every 5 min using HOBO Onset loggers (USA) to ensure accuracy of thermal profiles.

The maximum photosynthetic efficiency (F_v_/F_m_) of photosystem II of the algal symbionts was measured for each anemone at T7 (i.e., immediately after heat stress) and T18 (i.e., following an overnight post-stress recovery period) after 1 h dark acclimation [2], using a MINI-PAM-II fluorometer (Walz, Germany) based on established methods [4]. Thermal tolerance thresholds were calculated as the mean temperature reducing F_v_/F_m_ to 50% of baseline (Effective Dose 50, or ED50) [5]. Statistical analyses and visualization used R v4.1.0 [6] with ggplot2 [7] and tidyverse [8] packages. Data were plotted as mean ± standard error (SE). To enable robust statistical analysis, we only compared the ED50 thermal tolerance thresholds of Aipatsia H2 between the bacterial-inoculated group and its placebo control counterpart derived from the same CBASS run. The normality of the data distribution was assessed using the Shapiro-Wilk test, and the homogeneity of variances was evaluated with the F-test. For comparisons between the two groups (i.e., Control vs. MC10-B4, Control vs. MC0-A5, Control vs. MC15-BG7), an independent Student's t-test was employed. A *P*-value below 0.05 (*), 0.01 (**), and 0.001 (**) was considered statistically significant.

**KEGG module enrichment analysis**

To identify genomic features distinguishing MC10 members (n = 15) from those belonging to the other *Ruegeria* populations (n = 20), we first clustered all protein sequences predicted by Prokka [9] into orthologous groups (OGs) using OrthoFinder v2.5.1 [10]. Then, we selected the MC10-specific OGs based on the distribution of OGs across the 35 isolates. Thresholds were determined by visualizing the relationship between two metrics (see Figure 1A): the relative presence in MC10 (proportion of MC10 strains within an OG) and the overall presence across all strains (proportion of total strains harboring the OG). Based on the distribution of OGs in the resulting scatter plot, a 65% cutoff was identified as the optimal threshold where the two metrics intersected. Accordingly, MC10-specific present OGs (n=1,610) were defined by relative persistence in MC10 >= 65% and persistence in all strains < 65%. This ensured the selection of genes concentrated in MC10 while excluding universal core genes. MC10-specific absent OGs (n=4,873) were defined as those entirely missing in MC10 (0% occupancy) but present in the non-MC10 group. All individual protein sequences belonging to these specific OGs were then extracted as the target gene sets for functional investigation.

To ensure the most up-to-date biological interpretation, KEGG Module information was retrieved directly via the Biopython REST API (interfacing with KEGG Release 117.0+/02-12, Feb 26 Kanehisa Laboratories). A customized gene-set library was constructed by mapping the annotated KO (KEGG Orthology) identifiers to their corresponding KEGG Modules, representing well-defined functional units and metabolic segments. KO identifiers without any module information were removed before conducting the following enrichment analysis. To elucidate the biological significance of the MC10-specific genomic features, over-representation analysis (ORA) was conducted at the gene level using the enrich function from the Python package GSEApy package v1.1.11 [11]. For each enrichment run, the target gene set consisted of all protein members within the MC10-specific present or MC10-specific absent OGs. The enrichment was assessed using a hypergeometric test against a background (universe) of all annotated proteins from the 35 bacterial genomes. Modules with a p-value < 0.05 were defined as significantly enriched.

To account for phylogenetic autocorrelation, Phylogenetic Generalized Least Squares (PGLS) was employed using the caper package (v1.0.4) in R. This analysis validated whether the observed functional enrichments were significantly driven by the MC10 members identity beyond the effects of shared ancestry. The variance-covariance (VCV) matrix was derived from the pruned 35-strain phylogenomic tree and Pagel's λ parameter was estimated by maximum likelihood. The high-confidence modules (GSEApy p-value < 0.05 and PGLS nominal p-value < 0.05) were visualized using bubble plots.

**Phylogenomic Evaluation of KO Distributions**

For each KO, phylogenetically controlled statistics were performed using binary Phylogenetic Generalized Linear Mixed Model (binaryPGLMM) via the R package ape (v5.8.1). This approach accounts for phylogenetic non-independence by incorporating a tree-derived variance-covariance matrix, allowing for a robust evaluation of clade-specific patterns.

***Ruegeria* morphological analysis**

To investigate the unique functional adaptations of MC10, we first conducted a comparative morphological analysis. We focused on the representative strain MC10-B4, comparing it against two other coral-associated strains, MC0-A5 and MC15-BG7, which were selected from lineages that co-occur with MC10 and share the same sampling site, coral host species, individual colony, and tissue compartment. The model pelagic bacterium *R. pomeroyi* DSS-3 was included to provide a broader phylogenetic context.

        For electron microscopic analysis, we followed the established procedures for sample preparation [12]. Pure *Ruegeria* isolates were cultured in 3 mL of Marine Broth (MB) 2216 (BD Difco, USA) with shaking at 280 rpm at 28 °C for approximately 30 hours to reach the exponential growth phase. Cells from four *Ruegeria* strains were each harvested by centrifugation at 8,000 × g for 5 min at 4 °C for 5 min. The cell pellets were then washed twice with 1X PBS and resuspended in 1 mL of PBS. A 2% ultra-low gelling temperature agarose solution (Sigma, USA) was prepared by dissolving agarose powder in double-distilled water (ddH2O) with gentle heating. This solution was mixed with bacterial suspensions at a 1:3 (v/v) ratio to achieve a final agarose concentration of 0.5%. The mixture was maintained at 28 °C to prevent solidification prior to High Pressure Freezing (HPF). For HPF processing, 20 µL aliquots of the cell-agarose mixture were loaded into Type B standard HPF carriers (Science Services, Germany). Frozen samples were transferred under liquid nitrogen to 1 mL of anhydrous acetone containing 2% osmium tetroxide, followed by Autonomic Freeze Substitution (AFS) in a portable cryogenic refrigerator (CRYO PORTER, Model CS-80CP, Scinics Corporation, Japan), starting at -80 °C for 54 hours, during which pre-cooled 0.1% uranyl acetate was added to enhance contrast. The temperature gradually increased to -20 °C over a 24-hour period, and then gradually increased from -20 °C to -4 °C over 12 hours, followed by a further gradual warming to 4 °C over the course of 96 hours. Afterward, samples were washed four times with 100% acetone for 15 min each to eliminate any residual fixatives. The samples were then embedded in Epon resin (EMbed 812 Kit, Electron Microscopy Sciences, USA) by sequentially increasing acetone concentration (10%, 25%, 50%, and 75%), with each concentration maintained for at least 4 hours. Subsequently, the samples were infiltrated with pure Epon resin overnight, followed by an additional 8 hours of infiltration with freshly supplied pure Epon resin. Finally, the samples were incubated in pure degassed Epon resin supplemented with 2.5% DMP-30 shortly before being poured into molds for polymerization at 60 °C for 48 hours. After sample mounting, sections approximately 80 nm in thickness were cut using an ultramicrotome (Leica UC7, Germany) and collected onto nickel grids. Three sections for each sample were examined by transmission electron microscope (Hitachi H-7650, Japan).

**Motility of *Ruegeria* strains**

Swimming and swarming motility assays were performed as previously described with minor modifications [13]. Briefly, exponential-phase cultures of the four *Ruegeria* strains (i.e., MC10-B4, MC15-BG7, MC0-A5, and DSS-3) were adjusted to an OD_600_ of 0.5 in fresh MB. Aliquots (5 µL) of each bacterial suspension were spot-inoculated onto the surface of swimming plates (MB containing 0.18% agar) and swarming plates (MB containing 0.75% agar). The inoculated plates were incubated at 28 °C for 6 days (swimming motility) or 10 days (swarming motility). After incubation, the motility halos were photographed, and their diameters were measured. For each halo, both the length and width were recorded, and the average diameter was calculated. All experiments were performed in three independent biological replicates.

        For the twitching motility assay, 1 µL of the above bacterial suspension (OD_600_ = 0.5) was stab-inoculated into the bottom of 1% MB agar plates. All plates were incubated at 28 °C for 10 days. Post-incubation, the agar layer was carefully removed. The exposed adherent cells were dried at 55 °C for 10 min. The fixed cells were then stained with 0.1% (w/v) crystal violet for 15 minutes. Following staining, unbound dye was rinsed off with distilled water, and the plates were air-dried prior to imaging. The experiment was conducted with three independent biological replicates, and representative images are shown.

**Sedimentation assays**

The sedimentation assay was adapted from a previous study [13]. Exponential-phase cultures (OD_600_ = 0.5) of the four *Ruegeria* strains (i.e., MC10-B4, MC15-BG7, MC0-A5, and DSS-3) were diluted 1:1 in fresh MB. A 2 mL aliquot of each diluted suspension was placed in a 14-mL tube and incubated without shaking at room temperature. The sedimentation status was visually assessed and photographed at 24-hour and 48-hour time points. Three independent biological replicates were performed for every strain under identical conditions.

**Biofilm production of *Ruegeria* strains**

Biofilm formation was quantified using a crystal violet-based assay adapted from Smith et al. [14]. Briefly, bacterial cultures of the four *Ruegeria* strains (i.e., MC10-B4, MC15-BG7, MC0-A5, and DSS-3) in the exponential phase were adjusted to an OD_600_ of 0.1 in fresh MB. Aliquots of 150 µL were dispensed into the perimeter wells of a sterile 96-well flat-bottom plate to enable clear visualization of both wall- and bottom-adhered biofilms, while the four corner wells were filled with MB as negative controls to minimize edge effects. After 24 and 48 h, planktonic cells were removed by gently rinsing the wells with PBS, and the plates were dried at 55 °C for 10 min. Biofilms were stained with 200 μL of 0.1% (w/v) crystal violet for 15 min, rinsed three times with PBS, and dried for 10 min. Subsequently, images were taken, and 0.2 ml of decolorizing solution [methanol (4): acetic acid (1): water (5)] was added to each well, followed by incubation with gentle shaking at room temperature for 10 min. The absorbance of the resulting solution was measured at 590 nm. The final OD_590_ value for biofilm formation was obtained by subtracting the absorbance of the blank control (i.e., MB without bacteria processed identically). Experiments were performed with six biological replicates for each condition, and all test strains were analyzed simultaneously on the same plate.

**Siderophore production of *Ruegeria* strains**

Siderophore synthesis was assessed using the traditional Chrome Azurol S (CAS) assay, which is based on the competition between siderophores and the CAS-iron complex, resulting in a color change from blue to orange when siderophores are present [15]. The base MM9 medium was prepared by dissolving 0.9 g of KH_2_PO_4_, 1.5 g of NaCl, and 3 g of NH_4_Cl in 255 mL of ddH_2_O, with the pH adjusted to 6.5. Subsequently, 9.7 g of PIPES buffer and 4.5 g of agar were added to the solution. The blue dye solution was prepared by dissolving 0.06 g of CAS in 50 mL of ddH_2_O, 0.0027 g of FeCl_3_•6H_2_O in 10 mL of 10 mM HCl, and 0.073 g of hexadecyltrimethylammonium bromide (HDTMA) in 40 mL of ddH_2_O; the three solutions were then combined and mixed thoroughly. The MM9 medium and the blue dye solution were autoclaved separately, and then cooled to 50 °C. The components were then thoroughly mixed by combining 255 mL of MM9 with 33 mL of the blue dye solution, 10 mL of 10% casamino acids, and 3.3 mL of 20% glucose (that had been both filter-sterilized using a 0.22 µm membrane filter) to create the final solution. This solution was then used for making CAS agar plates. Single colonies of MC10-B4 and its counterpart strains (i.e., MC15-BG7, MC0-A5, and DSS-3) pre-cultured to the exponential growth phase were picked and then streaked onto the CAS agar plates, which were then incubated at 28 °C for 3 days. After incubation, plates were examined for the color change from blue to orange in the agar surrounding colonies to determine the production of siderophores.

**Vitamin prototrophy assay of *Ruegeria* strains**

Genome analysis predicted that strain MC10-B4 is unable to synthesize vitamin B_7_ (biotin) and vitamin B_12_ (cobalamin). To validate these predictions, cultures of the four *Ruegeria* strains (i.e., MC10-B4, MC15-BG7, MC0-A5, and DSS-3) were grown in a marine basal medium (MBM) under four vitamin supplementation conditions: (i) a complete vitamin supplement (following the recipe for medium 197 of the Japan Collection of Microorganisms [16]), (ii) no vitamin, (iii) all vitamins except B_7_, and (iv) all vitamins except B_12_. The MBM base, prepared as described previously [17], was used to control other nutrients and minimize confounding factors. Glucose was provided as the sole carbon source at a final concentration of 11 mM. Pre-cultured strains at the exponential growth phase were diluted to an OD_600_ of 0.1 and inoculated into the respective MBM with its specific vitamin supplementation. Cultures were incubated at 28 °C and 200 rpm for 144 hours, and growth curves were monitored over the course time by measuring the OD_600._

**Oxidative stress sensitivity assay**

The oxidative stress sensitivity of the strains was assessed using two complementary methods as described previously [13, 18] with minor adaptations. Briefly, exponential-phase cultures of strains MC10-B4, MC15-BG7, MC0-A5, and DSS-3 were harvested and adjusted to an OD_600_ of 0.5 in fresh MB. For the spot assay, the cell suspensions were then serially diluted (10-fold) in sterile normal saline (0.9% NaCl). Aliquots (5 μL) of dilutions from 10^-2^ to 10^-7^ were spotted onto MB agar plates supplemented with 0-, 0.05-, or 0.1-mM hydrogen peroxide (H_2_O_2_). Alternatively, for the liquid kill assay, the adjusted cell suspensions were incubated with or without 1 mM H_2_O_2_ at 28 °C for 1 hour. After treatment, the cells were serially diluted and spread on MB agar plates. For both assays, the MB agar plates were incubated at 28 °C for 3 days. The plates were photographed, and bacterial survival was quantified by counting the colony-forming units (CFU).

**Catalase activity assay**

Catalase activity was determined by the slide method with minor modifications [19]. Briefly, fresh bacterial colonies were transferred and smeared onto a glass slide. A 10 µL drop of 3% H_2_O_2_ was added, and the mixture was observed immediately. The production of visible oxygen bubbles indicated the presence of catalase enzymes.

**Coral tissue extract preparation**

Five coral mother colonies of *Acropora pruinosa* (Brook, 1893) were sampled from Crescent Island in Hong Kong waters (22°31'26.5"N, 114°19'03.6"E) at depths ranging from 1.2 m to 2.9 m in May 2024. Each mother colony contained 3-5 branches, each 3-4 cm in length. Coral fragments were transported in the cool box back to the Marine Science Laboratory at the Chinese University of Hong Kong within an hour and processed immediately upon arrival.

        Coral tissue extract (CTE) was prepared following previously described methodologies [20]. The tissue of each coral fragment was removed with 150 mL autoclaved and 0.22 μm-filtered seawater (AFSW) using WaterPik (WaterPik, US). The tissue suspension was centrifuged at 4 °C, 3,000 × g for 5 min, and the pellet was resuspended in 10 mL AFSW. Aliquots of 2 mL were distributed into 5 mL tubes, and homogenized on ice at 3,600 rpm for 30 s using a digital homogenizer T-18 ULTRA-TURRAX (IKA, Germany). The homogenate was centrifuged at 4 °C, 3,500 × g for 4 min to pellet the symbiont algal cells. Supernatant was carefully collected to avoid disturbing the pellet, then filtered through a 0.45 µm cell strainer to remove large particles. The filtrate was then loaded into a 3 kDa Amicon® Ultra Centrifugal Filter (Merck, USA), centrifuged at 4 °C, 3,000 × g for 3 hours, and the fraction with a molecular weight smaller than 3 kDa was collected. The fractionated CTE was pooled and stored at -80 °C for less than one week before the experiment.

**Bacterial incubation experiment**

Pure cultures of MC10-B4 and its coral-derived counterpart strains (i.e., MC15-BG7 and MC0-A5) were incubated at 28 °C with shaking at 280 rpm to reach their exponential phase. A 1 ml aliquot of each *Ruegeria* culture was subsequently transferred to 150 mL of MB and incubated for 16 hours at 28 °C with shaking at 280 rpm. After incubation, cultures were centrifuged at 8,000 × g at 4 °C for 5 min to pellet the bacterial cells. The pellets were then washed twice with AFSW and resuspended either in 50 mL of AFSW (control group) or in 45 mL of AFSW supplemented with 5 mL CTE (CTE-treated group). The 50 mL bacterial suspensions were dispensed into 8 mL aliquots in 15 mL tubes and incubated at 28 °C with shaking at 200 rpm for 4 hours. After treatments, *Ruegeria* cultures were centrifuged at 4 °C at 8,000 × g for 5 min, and pellets were washed twice with 1X PBS. The supernatant was discarded, and bacterial pellets were snap-frozen in liquid nitrogen and stored at -80 °C for subsequent protein extraction.

**Extraction, digestion, and sequencing of proteins from *Ruegeria* cells**

For protein extraction, frozen cell pellets of *Ruegeria* cultures were resuspended in 100 µL of 1X PBS and mixed with 25 µL of 5X SDS loading buffer. The mixture was incubated at 99 °C for 10 min to facilitate cell lysis. Lysates were centrifuged at 15,000 × g for 30 seconds, and 10 µL of the supernatant was used for SDS-PAGE protein separation. Following electrophoresis, the gel was stained with Coomassie Brilliant Blue for 10 min with gentle agitation, followed by overnight destaining with Milli-Q water. Protein bands were excised from the gel, cut into small pieces, and washed with 50% methanol and 50 mM NH_4_HCO_3_ until the blue color disappeared. The gel pieces were cleaned with 100% acetonitrile (ACN) and subsequently rehydrated in 10 mM NH_4_HCO_3_.

        Disulfide bonds were reduced by incubating the samples with 10 mM 1,4-dithiothreitol (DTT) in 10 mM NH_4_HCO_3_ at 56 °C for 40 min. Alkylation was performed by incubating with 55 mM iodoacetamide (IAA) in 10 mM NH_4_HCO_3_ at 24 °C in the dark for 20 min. Excess reagents and reaction byproducts were removed using 100% ACN, and the gel was rehydrated in 10 mM NH_4_HCO_3_. In-gel digestion was performed overnight at 37 °C using sequencing-grade trypsin (Promega, USA) with a concentration of 0.8 ng/µL. Peptides were extracted using a solution of 5% formic acid in 50% ACN for 15 min, then concentrated to 10-20 µL using SpeedVac. Finally, all samples were desalted using Pierce™ C18 Spin Columns (Thermo Scientific, USA) according to the manufacturer’s instructions and dried before proteomic sequencing. Peptide concentration was determined using a BCA protein assay kit (Thermo Fisher Scientific, USA), and 300 ng of peptides per sample were reconstituted in 0.1% formic acid. The prepared samples were analyzed on an Orbitrap Fusion Lumos Tribrid Mass Spectrometer (Thermo Fisher Scientific, USA) using settings as described previously [21].

**Proteomic data processing and analysis**

For accurate protein identification, strain-specific reference databases were created from the annotated genomes of three *Ruegeria* strains (i.e., MC10-B4, MC0-A5, and MC15-BG7). Each sample was exclusively matched against the database of its corresponding strain [22]. Common contaminant proteins were appended to each database for filtration purposes [23]. Mass spectra were searched against the reference databases using Proteome Discoverer (v3.1.0) [24], following the standard processing workflow with Minora feature detection and the Sequest HT search engine. This platform was chosen based on literature demonstrating its superior performance in quantification yield, dynamic range, and reproducibility [25].

        A typical label-free quantification (LFQ) workflow was applied to normalize spectral intensity, accounting for variations in sample loading without labeling. The relative protein abundance was calculated by summing the intensity of all peptides assigned to each protein. The “Replicated Based Resampling” mode in the “Precursor Ions Quantifier node” was enabled for imputation of missing values. This method imputes missing values for low-abundance signals by modeling the relationship between variability and abundance across replicates, drawing random values from a normal distribution around the median detected value [26-29]. In the quantification node, we utilized the ‘Feature Mapper’ node to perform retention-time alignment and feature linking across raw files, including gap filling to identify undetected features, thereby increasing coverage and consistency in peptide and protein quantifications across different samples. After protein identification and quantification, contaminant proteins were removed. Only proteins identified by at least two distinct peptides with complete quantification data were retained. Additionally, proteins were clustered based on OG identification to avoid redundancy from multiple proteins in the same group [23, 30, 31].

        Beta diversity across samples was evaluated using Principal Coordinates Analysis (PCoA) based on Bray-Curtis distance matrices. Differential protein abundance of OG groups between CTE-treated and control samples was analyzed using a two-tailed t-test with the limma algorithm. Significantly differentially expressed OG groups (DE_OGs) were identified for each *Ruegeria* strain by applying a false discovery rate (FDR) threshold of < 0.05 and an absolute log_2_ fold change of |log_2_FC| > 0.5.
